# Broadly Conserved TMEM-131 Family Proteins in Collagen Production and Secretory Cargo Trafficking

**DOI:** 10.1101/794842

**Authors:** Zhe Zhang, Meirong Bai, Guilherme Oliveira Barbosa, Andrew Chen, Yuehua Wei, Shuo Luo, Xin Wang, Bingying Wang, Tatsuya Tsukui, Hao Li, Dean Sheppard, Thomas B. Kornberg, Dengke K. Ma

## Abstract

Collagen is the most abundant protein in animals. Its dysregulation contributes to ageing and human disorders including tissue fibrosis in major organs. How premature collagens in the endoplasmic reticulum (ER) assemble and route for secretion remains molecularly undefined. From an RNAi screen, we identified an uncharacterized *C. elegans* gene *tmem-131,* deficiency of which impairs collagen production and activates ER stress response. TMEM-131 N-termini contain bacterial PapD chaperone-like (PapD-L) domains essential for collagen assembly and secretion. Human TMEM131 binds to COL1A2 and TRAPPC8 via N-terminal PapD-L and C-terminal domain, respectively, to drive collagen production. We provide evidence that previously undescribed roles of TMEM131 in collagen recruitment and secretion are evolutionarily conserved in *C. elegans*, *Drosophila* and humans.

---

Collagen is the major extracellular component of connective tissues and is the most abundant protein in animals (*1–3*). Production of mature collagen is a multi-step process, involving collagen gene regulation, protein biosynthesis, post-translational modifications in the endoplasmic reticulum (ER), formation of secretion-competent trimers, and extracellular C-propeptide cleavage and cross-linking among trimers (*1–3*). Dysregulation of collagen production or deposition contributes to a wide variety of human disorders, including diabetes, ageing and pathological tissue fibrosis in major organs such as kidneys, liver, lungs and heart (*4–9*). The type I collagen is the most abundant fibril-type collagen and its trimer comprises two α1 (I) procollagen chains and one α2 (I) procollagen chain, encoded by the genes *COL1A1* and *COL1A2*, respectively. Both COL1A1 and COL1A2 contain C-terminal domains (C-propeptide) responsible for initial chain trimerization. The trimerization occurs via a zipper-like mechanisms, initiating from the C-propeptide domain of α1/2 (I) in close proximity to ER membranes (*10–13*). Although the enzymes responsible for type I collagen modification, extracellular cleavage and cross-linking have been well described (*1, 14*), how procollagen monomers are recruited to assemble into secretion-competent multimers in the ER remains poorly understood.

COPII-coated vesicles mediate ER-to-Golgi anterograde transport of secretion-competent cargos, including those containing collagens (*15, 16*). Tethering COPII vesicles from ER to Golgi membranes requires TRAPP (transport protein particle), a multi-subunit protein complex highly conserved in eukaryotes (*17, 18*). TRAPPC8 is a key component of TRAPP III, a subtype of TRAPP that acts as a guanine nucleotide exchange factor (GEF) to activate Rab GTPase to promote ER-to-Golgi cargo trafficking. Collagen production also requires HSP47, a procollagen chaperone in the ER, and TANGO1, an ER transmembrane protein that facilitates the export of bulky cargo with collagen (*19, 20*). Intriguingly, HSP47 and TANGO1 orthologs are present only in vertebrates (*21–23*), raising the question whether mechanisms conserved in all animals exist to recognize procollagens for assembly into export-competent collagen trimers in route for secretion.

In this study, we identify a previously uncharacterized *C. elegans* gene *tmem-131* that defines an evolutionarily conserved protein family important for procollagen recruitment and secretion. The exoskeleton of *C. elegans*, is a complex collagen matrix that comprises many distinct mature collagen proteins, including COL-19, an adult-specific, hypodermally synthesized collagen (*24*). From yeast-two-hybrid (Y2H) screens, we identified two human proteins COL1A2 and TRAPPC8 that bind to the N- and C-terminal domains of hTMEM131, respectively, and show that COL-19 secretion requires TMEM-131 and TRPP-8, the *C. elegans* ortholog of TRAPPC8. TMEM131 proteins are also essential for collagen secretion in *Drosophila* and human cells, supporting evolutionarily conserved role of TMEM131 protein family in collagen production.

## Genome-wide RNAi screen identifies *tmem-131* regulating ER stress response and collagen production in *C. elegans*

We performed a genome-wide RNAi screen for genes that affect levels of the *C. elegans* transgenic reporter *asp-17*p::GFP (Fig. 1A). *asp-17* is a gene encoding an aspartyl protease-like protein that is up-regulated by temperature stress and down-regulated by ER stress (Fig. 1A-C) (*25*). From a screen of over 19,100 genes, we identified 574 RNAi clones that either up-or down-regulated the abundance of *asp-17*p::GFP (Table S1). In this work, we focus on the gene *tmem-131*, as it is uncharacterized but otherwise highly evolutionarily conserved in all animals (see below). RNAi against *tmem-131* caused a fully penetrant and strong suppression of *asp-17*p::GFP (Fig. 1B). By contrast, RNAi against *tmem-131* caused marked up-regulation of *hsp-4*p::GFP (Fig. 1C), an established transcriptional reporter for ER stress and unfolded protein response (UPR) in *C. elegans* (*26–28*). The screen also identified many other genes, including *ostb-1* and *dlst-1* that are involved in protein modification and homeostasis in the ER (Fig. 1C). RNAi against *ostb-1* or *dlst-1*, as *tmem-131*, also caused marked up-regulation of *hsp-4*p::GFP and down-regulation of *asp-17*p::GFP (Fig. 1C).

**Fig. 1.**
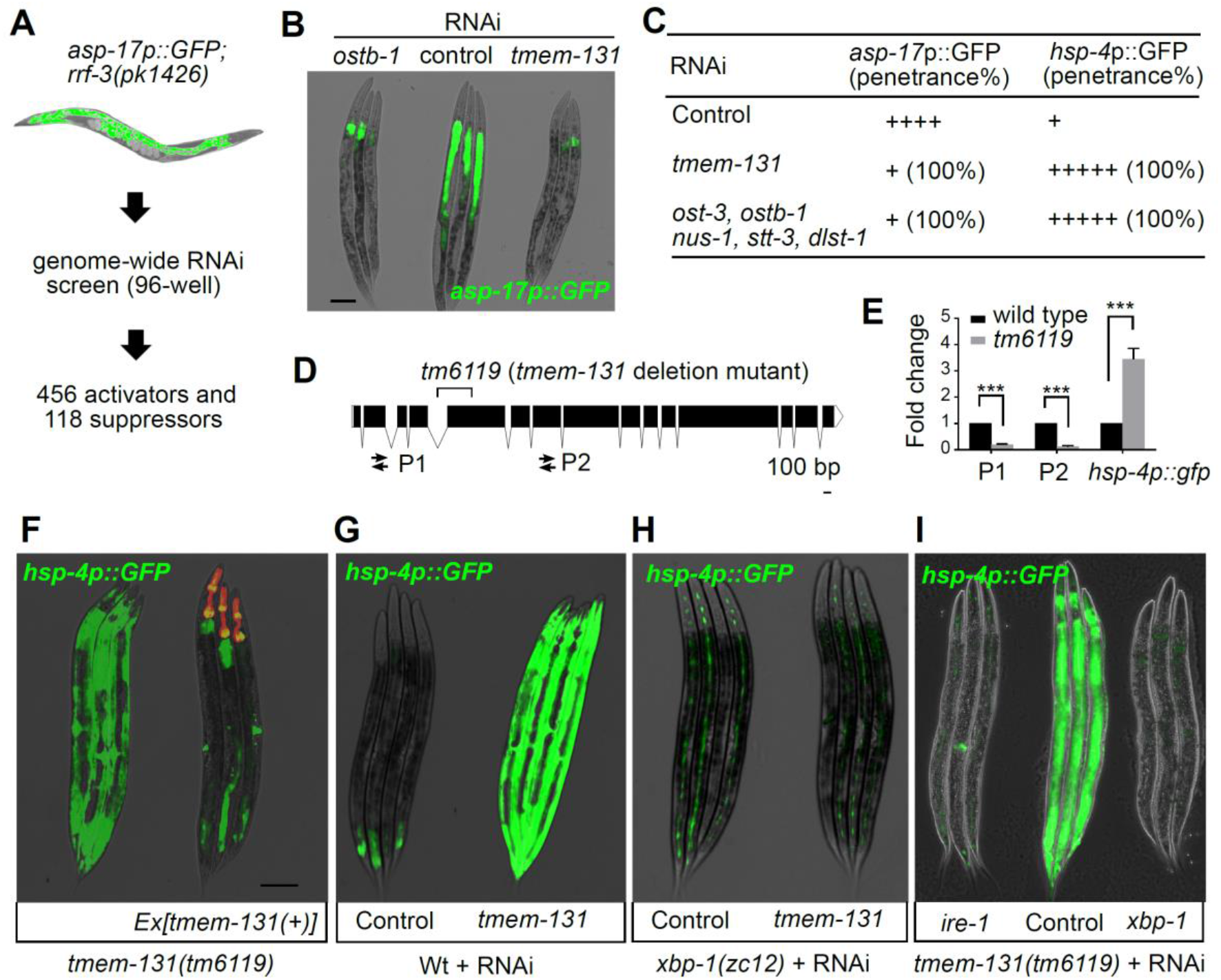
Genome-wide RNAi screen identifies *tmem-131* regulating ER stress response in *C. elegans*. **(A)**, Schematic of RNAi screen for regulators of *asp-17p*::GFP. (**B)**, Exemplar fluorescence and bright-field images for *rrf-3; asp-17p*::GFP with control, *ostb-1* and *tmem-131* RNAi. **(C)**, Table listing ER proteostasis genes whose RNAi also suppress *rrf-3; asp-17p*::GFP. (**D)**, Schematic of Ce-*tmem-131* LOF alleles, and RT-PCR P1 and P2 primers for *tmem-131* mRNA measurement. (**E)**, Quantification of fold changes in *tmem-131* and *hsp-4::GFP* expression levels in wild type and *tm6119* mutants. *** indicates P < 0.001 (n ≥ 3 biological replicates). (**F)**, Exemplar fluorescence and bright-field images showing rescue of *hsp-4p*::GFP induction in *tm6119* mutants with transgenic expression of *tmem-131*p::*tmem-131*::*gfp* marked by pharyngeal *myo-2*p::mCherry. (**G-I)**, Exemplar fluorescence and bright-field images for the UPR reporter *hsp-4p*::GFP with *tmem-131* RNAi in wild type (**G**) and *xbp-1* mutants (**H**), and (**I**) in *tm6119* mutants with control, *xbp-1* or *ire-1* RNAi. Scale bars: 100 µm.

To verify the RNAi phenotype, we examined *C. elegans* mutants carrying a genetic deletion allele *tm6119*, which caused a protein-coding frame shift and severe reduction of *tmem-131* expression level (Fig. 1D, 1E). *tm6119* mutants exhibited abnormally elevated levels of *hsp-4*p::GFP that can be rescued by transgenic expression of wild-type *tmem-131* (Fig. 1F). In addition, high *hsp-4*p::GFP levels in *tm6119* mutants or *tmem-131* RNAi-treated animals were completely suppressed by loss of function (LOF) of XBP-1 (Fig. 1G-I), a transcription factor that drives a major branch of UPR in *C. elegans* (*27–30*). Loss of IRE-1, an ER-transmembrane protein that senses ER stress, also prevented *hsp-4*p::GFP up-regulation in *tm6119* mutants (Fig. 1I). Besides constitutively activated *hsp-4*p::GFP expression, we found that TMEM-131 deficient animals by RNAi or *tm6119* were smaller in size at higher temperature, more sensitive to the ER stressor tunicamycin as well as cuticle-disrupting osmotic stresses, and developed more slowly compared with wild type (Fig. S1). Nonetheless, unlike *hsp-4*p::GFP, these additional phenotypes were not suppressed by loss of *xbp-1* or *ire-1* (Fig. S1G). Together, these results indicate that loss of *tmem-131* causes various organismic phenotypes and defective ER homeostasis, leading to IRE-1 and XBP-1-dependent activation of *hsp-4*p::GFP and UPR.

To investigate the mechanism by which TMEM-131 may regulate ER function and proteostasis, we first examined its expression pattern and subcellular localization. The promoter of *tmem-131* drives expression of GFP in a variety of *C. elegans* tissues, most prominently the intestine and hypoderm (Fig. 2A). A translational reporter with GFP fused to the C-terminus of TMEM-131 driven by the endogenous *tmem-131* promoter reveals an intracellular perinuclear reticulum pattern (Fig. 2B). The translational reporter rescued the *tm6119* mutant phenotype, indicating the reticulum-localizing TMEM-131 is functional (Fig. 1F). In addition, SignalP-4.1 predicts an ER signal peptide sequence (a.a. 1-30) of TMEM-131, supporting its ER-endosomal localization (*31*). Its cellular loci and regulation of *hsp-4*p::GFP led us to hypothesize that TMEM-131 normally acts to regulate processing, trafficking and/or homeostasis of ER-resident proteins. To identify potential client proteins of TMEM-131, we screened a panel of translational fluorescent reporters for ER-resident transmembrane and secreted proteins, seeking any phenotypic defects caused by RNAi against *tmem-131* (Table S2, Fig. S2). Among 34 various reporters we comprehensively examined, the COL-19::GFP reporter displayed the most striking defect in GFP patterns caused by LOF of *tmem-131*.

**Fig. 2.**
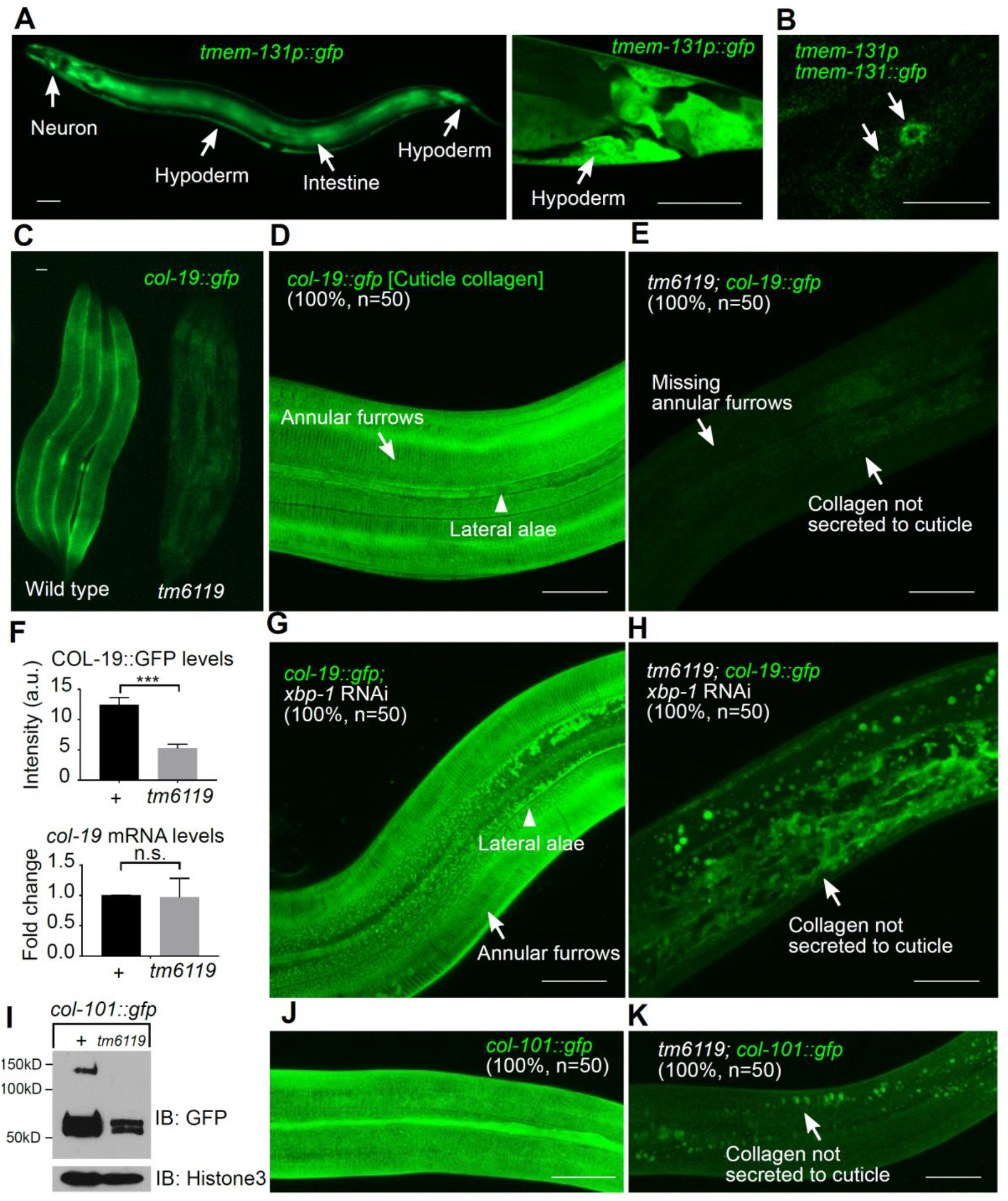
Ce-TMEM131 is essential for secretion of GFP-labelled collagen COL-19 and COL-101. **(A)**, Exemplar epifluorescence image of *tmem-131*p::GFP transcriptional reporter. (**B)**, Exemplar confocal image of *tmem-131p::tmem-131*::GFP showing vesicular puncta pattern in perinuclear areas of hypodermal cells. (**C)**, Exemplar epifluorescence image of *col-19*::GFP (i.e. *col-19*p::*col-19*::GFP) and *tm6119*; *col-19*::GFP. (**D-E)**, Exemplar confocal fluorescence images with indicated phenotypic penetrance of *col-19*::GFP in wild type (**D**) and *tmem-131* mutant (**E**). (**F)**, Quantification of COL-19::GFP fluorescent intensity and endogenous *col-19* mRNA levels in wild type and *tmem-131*(*tm6119*) mutants. **(G-H)**, Exemplar confocal fluorescence images with indicated phenotypic penetrance of *col-19*::GFP with *xbp-1* RNAi in wild type (**G**) or *tmem-131* mutants (**H**). *** indicates P < 0.001 (n ≥ 3 biological replicates). n.s., no significant differences. (**I)**, Exemplar SDS-PAGE and Western blot analysis of *col-101*::GFP and *tm6119*; *col-101*::GFP proteins from total animal lysates. (**J-K)**, Exemplar confocal images with indicated phenotypic penetrance of *col-101*::GFP in wild type (**J**) and *tmem-131* mutants (**K**). Scale bars: 20 µm.

COL-19 is a *C. elegans* collagen protein that is secreted by the hypoderm and required for integral structure of the cuticle (*24, 32*). COL-19::GFP deposition is enriched in the wild-type adult animal, constituting regular cuticle structures of annular furrows and lateral alae (Fig. 2C, 2D). By contrast, cuticle COL-19::GFP is completely absent in the *tm6119* mutant, with apparently missing COL-19::GFP-marked annular furrows and weak intracellular COL-19::GFP in the hypoderm (Fig. 2E). *tm6119* decreased the abundance of COL-19::GFP proteins without affecting the promoter activity or mRNA abundance of *col-19* (Fig. 2F, S2G). The striking COL-19::GFP phenotype was not caused by UPR via activation of XBP-1, since *xbp-1* RNAi restored *hsp-4*p::GFP expression to normal levels but not cuticle COL-19::GFP in *tmem-131* mutants (Fig. 2G, 2H). LOF of *xbp-1* by RNAi in the wild type background caused excessive ER stresses but did not apparently affect COL-19::GFP in the cuticle (Fig. 2G). In the *tmem-131* mutant background, *xbp-1* RNAi caused more prominent intracellular accumulation of COL-19::GFP (Fig. 2H). By Western blot analysis, we found that *tm6119* decreased overall COL-19::GFP abundance irrespective of environmental temperature (Fig. S2H), although the developmental or *hsp-4*p::GFP phenotype of *tm6119* mutants worsened at 25 °C compared with 15 °C or 20 °C (Fig. S1). In addition, we found that another collagen reporter COL-101::GFP similarly required TMEM-131 for normal production (Fig. 2I-K). *tm6119* decreased COL-101::GFP abundance most prominently in higher molecular weight species (Fig. 2I). RNAi against *pdi-2*, which is essential for collagen folding and assembly (*33*), also partially decreased abundance of COL-19 (Fig. S3A, B). RNAi against *cup-2*, the Derlin orthologue essential for ER-associated degradation (ERAD) (*34*), caused synthetic lethality with *tm6119* (Fig. S3C, D), consistent with the observation that abnormally accumulated COL-19::GFP as ERAD substrate decreased in overall abundance in *tmem-131* mutants (Fig. 2). These results indicate that TMEM-131 is essential for mature collagen production and its deficiency causes ER stress and UPR through abnormal accumulation of secreted proteins, including at least collagen COL-19 and COL-101 in the ER.

## Evolutionarily conserved roles of TMEM131 family proteins for collagen production

TMEM-131 has not been previously characterized although it belongs to the highly evolutionarily conserved Pfam12371 (TMEM131_like) protein family (*35–37*). Its protein sequence is predicted by the program TOPCONS to contain two transmembrane segments, the first (i.e. signal peptide targeting to the ER) and second of which span a hydrophilic domain facing ER, endosomal lumen or extracellular space while the C-terminal part is predicted to localize in the cytosol (Fig. 3A) (*38*). We used structure homology modelling (SWISS-MODEL) (*39*) to search for proteins structurally similar to TMEM131 and identified many of those from bacteria that contain the PapD chaperone domain involved in the assembly and secretion of extracellular pilus components (Fig. S4) (*40, 41*). The PapD-like domain (PapD-L) in the N-terminus of TMEM-131 is predicted to be in the ER lumen (Fig. 3A), consistent with a putative role in procollagen processing. SWISS-MODEL also predicted a second PapD-L in the N-terminus of TMEM-131 albeit with lower similarity score. Structural comparisons further revealed that the *C. elegans* PapD-L, when modeled against one of the most similar structural homolog proteins from *Porphyromonas gingivalis* W83, indeed comprises a two-loped immunoglobulin fold characteristic of canonic bacterial PapD domains (Fig. S4C, E) (*40, 42, 43*).

**Fig. 3.**
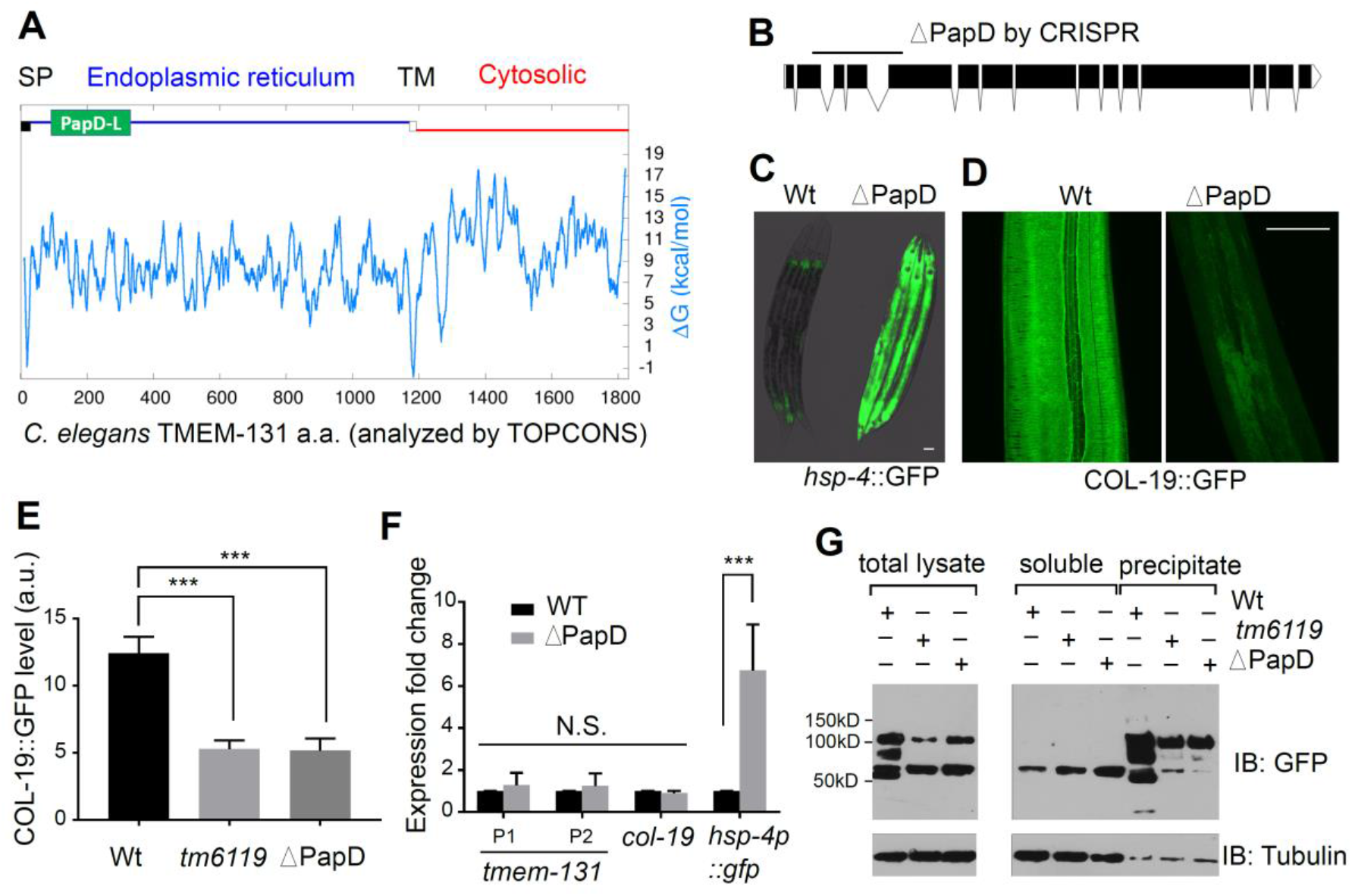
PapD-L chaperone domains are essential for TMEM-131 functions. **(A)**, Schematic of *C. elegans* TMEM-131 domain organization with PapD-L domain in green, ER localization in blue and cytosol localization in red as predicted by the TOPCONS program (http://topcons.net/). The biological hydrophobicity scale calculates the free energy (ΔG) of membrane insertion for a window of 21 amino acids (a.a.) centered on each position in the TMEM-131 sequence. (**B)**, Schematic of *tmem-131* gene structure with the PapD-L deletion generated by CRISPR/Cas9. (**C-D)**, Exemplar fluorescence images for *hsp-4p*::GFP (**c**) and *col-19*p::*col-19*::GFP (**D**) in wild type and PapD-L-deletion mutants. Scale bars: 20 µm. (**E)**, Quantification of the COL-19::GFP fluorescent intensity in wild type, *tmem-131*(*tm6119*) and *tmem-131*(ΔPapD-L) mutants. *** indicates P < 0.001 (n ≥ 3 biological replicates). (**F)**, Quantitative RT-PCR measurements of endogenous *tmem-131*, *col-19* and *hsp-4p::gfp* mRNA levels in wild type and ΔPapD mutants. *** indicates P < 0.001 (n ≥ 3 biological replicates). n.s., no significant differences. **G**, Exemplar Western blot analysis of COL-19::GFP in different fractions from wild type, *tmem-131*(*tm6119*) and *tmem-131*(ΔPapD-L) mutants.

To determine the physiological importance of PapD-L of TMEM-131, we generated a precise deletion of the coding sequence for the PapD-L domain in *C. elegans tmem-131* (Ce-PapD) using CRISPR/Cas9 (Fig. 3B). Ce-PapD deletion recapitulated the *tmem-131(tm6119)* or *tmem-131* RNAi phenotype in strong *hsp-4*p::GFP induction and defective COL-19::GFP secretion, although Ce-PapD deletion did not affect the overall abundance of *tmem-131* (Fig. 3C-F). The phenotypes caused by *tm6119* were rescued by the wild-type (100%, n = 20) but not the PapD-lacking (0%, n = 20) transgenes of *tmem-131*. We further analyzed COL-19::GFP proteins by Western blot analysis after separation of soluble versus insoluble fractions, which contain monomeric procollagens and multimerized/cross-linked collagens, respectively, from whole animal lysates. Consistent with the notion that secreted COL-19::GFP is cross-linked thus insoluble, we found that both *tm6119* and Ce-PapD deletion caused striking and robust decrease of the insoluble fractions of COL-19::GFP compared with wild type, while the abundance of soluble monomeric procollagens was less affected (Fig. 3G). These results indicate that Ce-PapD is essential for TMEM-131 to function in collagen recruitment and assembly, which is required for collagen secretion and preventing ER stress in *C. elegans*.

Within the evolutionarily conserved TMEM131_like protein family (Fig. 4A), the invertebrate model organisms *C. elegans* and *Drosophila* have one ortholog each, named *tmem-131* and *CG8370*, respectively. Vertebrate genomes encode two paralogs of TMEM-131, e.g. TMEM131 and KIAA0922 (a.k.a. TMEM131L) in humans (Fig. 4A, B). PapD-like domains are predicted from each homolog and represent the most highly conserved parts of TMEM131_like family proteins, based on multiple sequence alignment analysis (Clustal Omega) (Fig. S4A). We sought to identify conserved protein-interacting partners as the ligand or client proteins for Ce-PapD in TMEM-131 and used a yeast-two-hybrid (Y2H) screen to search for human proteins that can bind to Ce-PapD (Fig. 4C). From a normalized human cDNA library with approximately 9 million yeast clones in the screen, the Ce-PapD bait cDNA yielded 34 Ce-PapD interactor-encoding prey clones, among which COL1A2 was confirmed to interact with Ce-PapD in the Y2H assay (Fig. 4D, Table S3). COL1A2 constitutes the type I collagen fibril together with COL1A1 (*2, 44*). The library clone of human COL1A2 cDNA encodes the last 165 amino acid residues and a long 501 bp of the 3’ untranslated region of full-length COL1A2. We re-cloned the C-propeptide domain of COL1A2 and confirmed its specific interaction with Ce-PapD (Fig. 4D). In addition to Ce-PapD, we found that the PapD-like domains from the *Drosophila* TMEM131 homolog CG8370 and human TMEM131 but not TMEM131L (also known as KIAA0922) (*36*) proteins can also interact with the C-termini of human COL1A2 by Y2H assays (Fig. 4D). These results indicate that the TMEM131_like protein family members from *C. elegans*, *Drosophila* and humans exhibit evolutionarily conserved biochemical interactions with the type I collagen protein COL1A2 via their PapD-L domains.

**Fig. 4.**
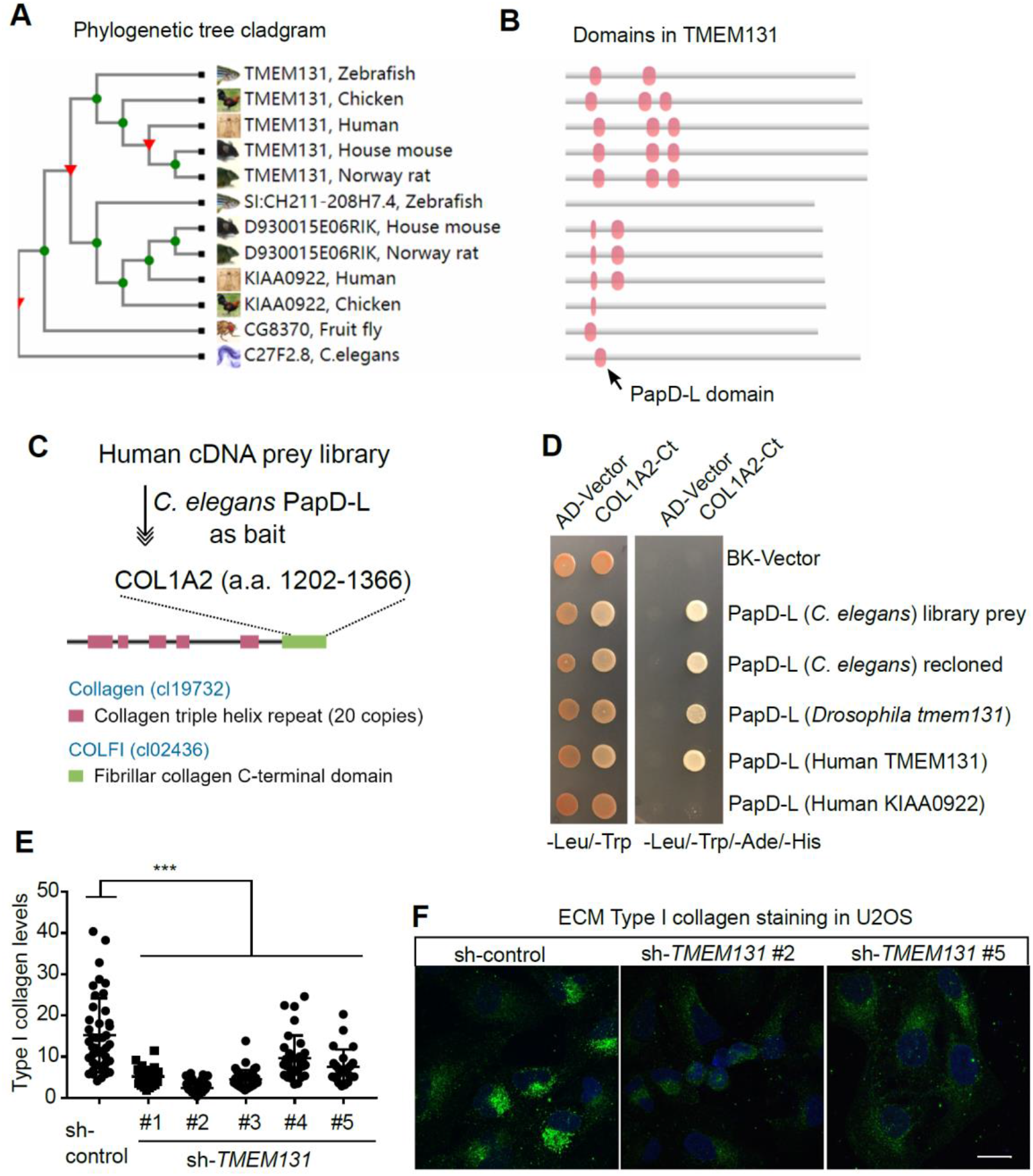
Evolutionarily conserved role of TMEM131 for collagen secretion in human cells. **(A)**, Cladgram of phylogenetic tree for the TMEM131 protein family from major metazoan species (adapted from Wormbase). (**B)**, Domain architectures of TMEM131 family proteins. (**C)**, Schematic of Ce-PapD-L Y2H screens identifying the human COL1A2 C-terminal domain as an evolutionarily conserved binder of PapD-L. (**D)**, Yeast colony growth in Y2H assays after re-transformation of prey and bait vectors verifying the interaction between human COL1A2 C-termini with PapD-L domains from *C. elegans*, *Drosophila* and humans. (**E)**, Quantitative immunofluorescence measurements of type I collagen abundance in control and *TMEM131* shRNA-depleted cells. *** indicates P < 0.001 (N > 10 cells for each condition; n ≥ 3 biological replicates). (**F)**, Exemplar confocal fluorescence images of U2OS cells with type I collagen staining after lentiviral expression of control shRNA and *TMEM131* shRNAs (#2 and #5). Scale bars: 20 µm.

To test whether TMEM131_like proteins might have evolutionarily conserved functions in collagen production, we examined LOF phenotype of *CG8370* and *TMEM131* in *Drosophila* and human cells, respectively. In human cells, we used lentiviral expression of small-hairpin RNAs (shRNA) to stably knock-down *TMEM131* in the collagen-producing bone osteosarcoma U2OS cell line (Fig. 4E, F, Fig. S5) (*45*). Immunofluorescence analysis revealed that *TMEM131* depletion markedly decreased abundance of type I collagens (Fig. 4E), the severity of which is largely correlated with the knockdown efficiency of shRNAs targeting 5 different coding sequences of *TMEM131* (Fig. 4E, F, Fig. S5). Decreased type I collagen in TMEM131-depleted human cells is consistent with diminished COL-19::GFP abundance of *C. elegans tmem-131* mutants (Fig. 2). Fully penetrant synthetic lethality of LOFs for both *tmem-131* and the Derlin gene *cup-2* strongly indicates that the ERAD pathway degrades procollagens if not properly assembled and secreted (Fig. S3D).

In *Drosophila*, we used the Lsp2>Collagen:RFP transgenic fly to visualize fat body-secreted collagen and RNAi to silence expression of *CG8370* in fat body cells (*46, 47*). We found that transgenic *CG8370* RNAi but not control animals accumulated Collagen::RFP in fat body cells, indicative of defective collagen secretion (Fig. 5). Interestingly, in *Drosophila* fat body cells in which collagen is normally secreted to the hemolymph (insect blood), *tmem131* LOF caused collagen accumulation but did not appear to involve further degradation by ERAD. Nonetheless, these results collectively provide evidence that roles of TMEM131 proteins are evolutionarily conserved for collagen secretion in at least human and *Drosophila* cells.

**Fig. 5.**
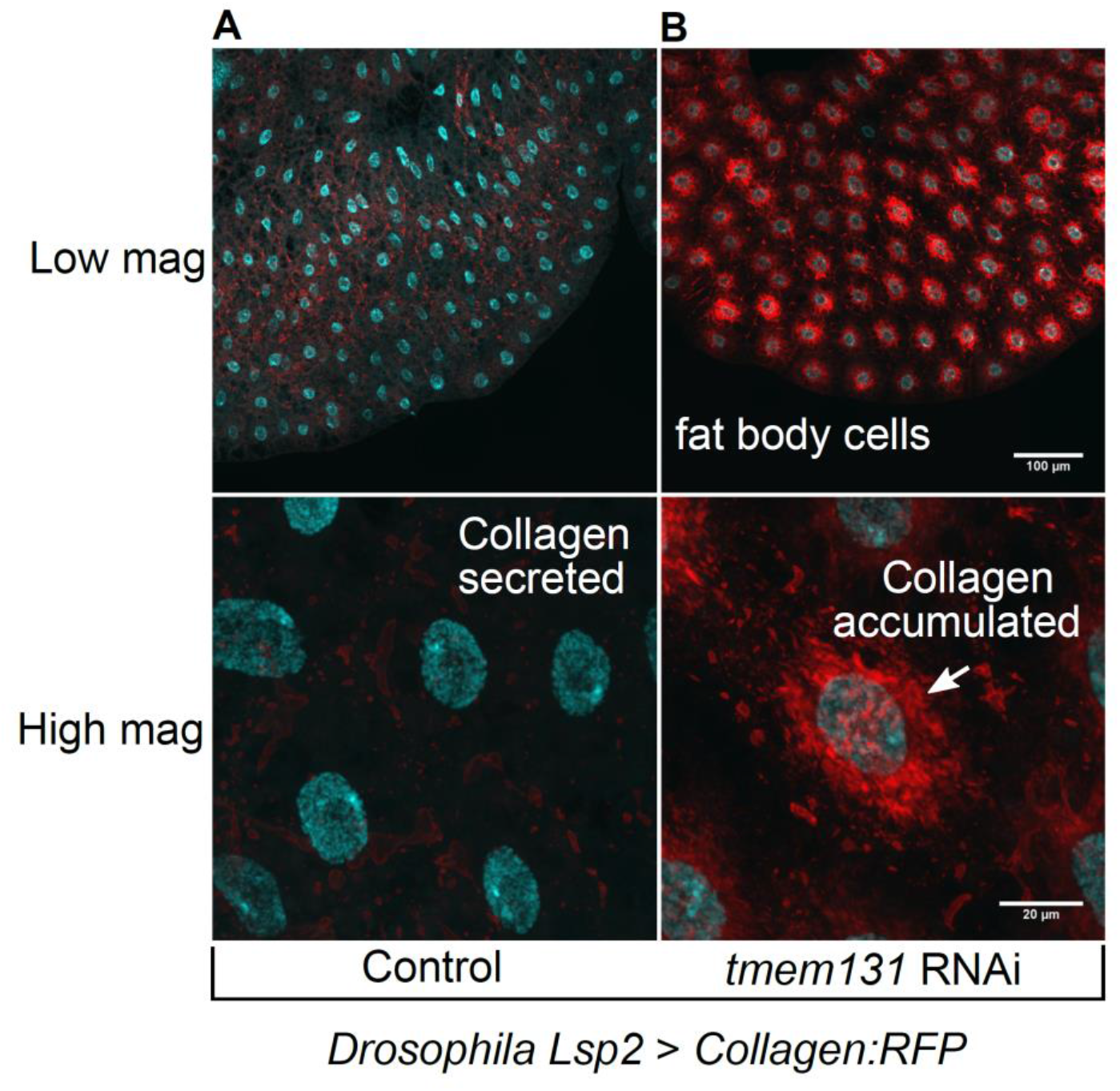
Evolutionarily conserved Role of TMEM131 for collagen secretion in *Drosophila*. **(A)**, Exemplar confocal images of *Drosophila* fat body cells showing normal collagen secretion with control transgenic RNAi. (**B)**, Exemplar confocal images of *Drosophila* fat body cells showing defective collagen secretion after fat body cell-specific transgenic RNAi of *Drosophila tmem131*. Shown are both low-mag (scale bar: 100 µm) and high-mag (scale bar: 200 µm) views indicating collagen accumulation in fat body cells because of defective secretion of collagen to the hemolymph (insect blood).

## TMEM131 promotes collagen cargo secretion through cytoplasmic C-terminal interaction with TRAPPC8

To address the mechanism by which the C-terminal domains of TMEM131 family proteins participate in collagen secretion, we used Y2H screens to identify proteins that can interact with the C terminus of human TMEM131 (Fig. 6A, Table S4). Among the prey cDNA clones identified to confer interaction with human TMEM131 Ct as the bait, we focused on TRAPPC8 in this study, as TRAPPC8-containing TRAPP III is critical for the ER-to-Golgi transport of collagen cargos. The TRAPPC8 prey clone identified encodes only part of its C-terminal region (Fig. 6A, B). We subsequently verified interaction of the re-cloned full-length TRAPPC8 with TMEM131 Ct using co-immunoprecipitation (CoIP) assays (Fig. 6C). When co-expressed in HEK293 cells, mCherry-tagged TMEM131 Ct specifically pulled down GFP-tagged TRAPPC8. Using Y2H assays, we found that substitution of a highly conserved tripartite motif WRD to AAA in TRAPPC8 Ct attenuated its interaction with TMEM131 Ct, whereas substitution of two highly conserved tryptophan residues to alanines in TMEM131 Ct attenuated its interaction with TRAPPC8 Ct, indicating that these residues are important for interaction and might constitute part of the interacting interface (Fig. 6D, Fig. S6). Consistent with interaction between TRAPPC8 and TMEM131, RNAi-mediated depletion of expression of *trpp-8,* the *C. elegans* homolog of *TRAPPC8*, strongly reduced the abundance of COL-19::GFP in the cuticle (Fig. 6E). RNAi against genes encoding additional components of TRAPP III also led to similar phenotype (Table S5). Together, these results indicate that TMEM131 promotes collagen secretion not only by its N-terminal PapD-L domain but also by its C-terminal recruitment of TRAPP III for the ER-to-Golgi transport of collagen cargo.

**Fig. 6.**
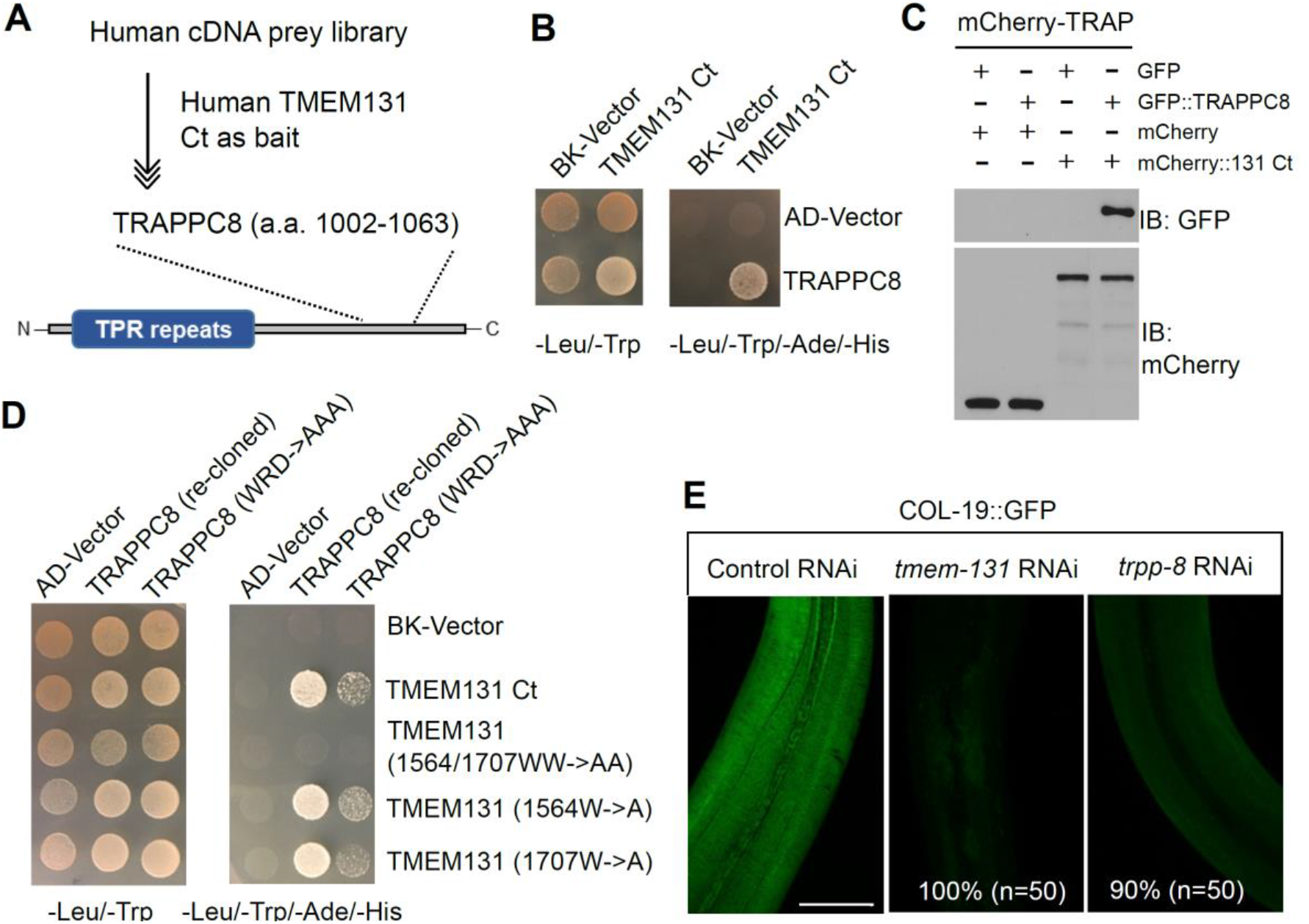
Roles of TRAPPC8 in binding to TMEM131 and collagen secretion. **(A)**, Schematic showing identification of TRAPPC8 as a TMEM131 interactor from Y2H screens. The C-terminus of human TMEM131 as a bait yielded a prey cDNA clone encoding the a.a. 1002 – 1063 C-terminal region of TRAPPC8. (**B)**, Yeast growth colonies verifying interaction of TMEM131 Ct and TRAPPC8 Ct. (**C)**, CoIP and Western blot showing biochemical interaction of GFP-labeled TRAPPC8 and mCherry-labelled TMEM131 Ct in HEK293 cells. Cells were co-transfected with expression vectors, lysed for immunoprecipitation by mCherry-TRAP, and blotted by antibodies against GFP and mCherry. (**D)**, Yeast growth colonies testing interaction of various TMEM131 Ct mutants with TRAPPC8 Ct or a mutant carrying WRD->AAA substitution, which attenuated interaction. (**E)**, Exemplar fluorescence images showing strongly decreased COL-19::GFP in *C. elegans* treated with RNAi against *tmem-131* or *trpp-8*. Scale bar: 50 µm.

Interaction of TMEM131 C-terminus with TRAPPC8 prompted us to determine the *in vivo* consequence of deleting *tmem-131* C-terminus on UPR and COL-19 in *C. elegans*. We used CRISPR-Cas9 to generate a precise deletion of the entire cytoplasmic domain of *tmem-131* and crossed the mutant to *hsp-4*p::GFP and COL-19::GFP reporters (Fig. 7A, B). We found that C-terminal deletion mutants showed strong activation of *hsp-4*p::GFP and reduction of COL-19::GFP abundance in cuticles. To identify specific sub-region of TMEM131 Ct responsible for interaction with TRAPPC8, we generated a series of TMEM131 Ct deletion mutants and test their interaction with TRAPPC8 in Y2H assays (Fig. 7C). Among the 5 deletions spanning the 1142 – 1883 a.a. sequence, only the very C-terminal end deletion (1741 – 1883 a.a.) abolished interaction with TRAPPC8 (Fig. 7D). We also generated 5 protein-coding mutations at evolutionarily conserved sites of TMEM131 Ct but found none of the single mutation can abolish the interaction with wild-type TRAPPC8 (Fig. 7E). By contrast, the C-terminal WRD->AAA mutation in TRAPPC8 attenuated interaction with wild-type TMEM131, indicating that the interaction interface between the two may require the WRD motif and multiple residues in TMEM131. Together, these results show that the C-terminal tail domain (named as TRAPID, for <u>TRAP</u>P III-Interacting <u>D</u>omain) of TMEM131 binds to TRAPPC8 (Fig. 7F) and its *C. elegans* counterpart is essential for collagen production *in vivo*.

**Fig. 7.**
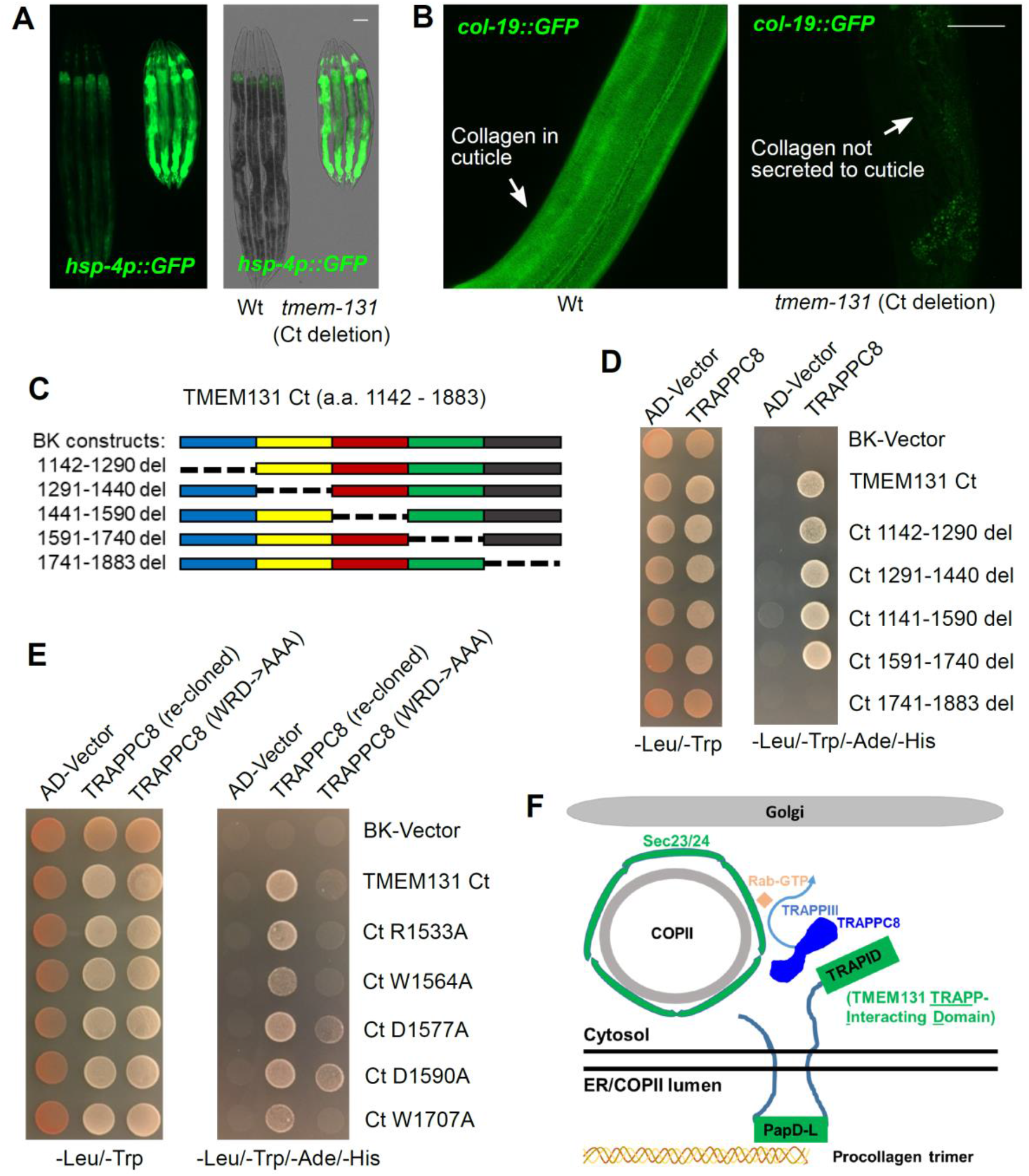
TMEM131 promotes collagen secretion through cytoplasmic C-terminal interaction with TRAPPC8. **(A)**, Exemplar fluorescence (left) and DIC merged (right) images showing activation of *hsp-4p::GFP* by CRISPR-mediated specific deletion of *C. elegans tmem-131* cytoplasmic C terminus. (**B)**, Exemplar fluorescence images showing the abolished COL-19::GFP signal in the cuticle of mutants (right) with deletion of *tmem-131* cytoplasmic C terminus compared with wild type (left). Scale bars: 50 µm. (**C)**, Schematic showing a series of deletions in human TMEM131 used for Y2H assays to identify regions responsible for binding to TRAPPC8. (**D)**, Yeast colonies from Y2H assays showing interaction of TRAPPC8 with various TMEM131 Ct mutants. (**E)**, Yeast growth colonies from Y2H assays showing interaction of TRAPPC8 and a WRD->AAA mutant with TMEM131 Ct mutants with single alanine substitutions in the conserved C-terminal domain. (**F),** Schematic model illustrating how TMEM131 regulates collagen secretion through its N-terminal PapD-L domain and C-terminal TRAPID domain (TMEM131 <u>TRAP</u>P-Interacting <u>D</u>omain). PapD-L binds to the C-propeptide domain of COL1A2 and facilitates assembly of procollagen trimers. TRAPID binds to TRAPPC8 and facilitates TRAPP III activation of Rab GTPase to promote the ER-to-Golgi transport of collagen cargo in COPII. For clarity, procollagen trimers in COPII and other essential factors for collagen secretion, including TANGO, HSP47 and SEC13/31 are not shown.

## Discussion

In this work, we identify a *C. elegans* protein TMEM-131, which has homologs in most animals, with essential roles in collagen production. Such roles appear evolutionarily conserved for *Drosophila* and human TMEM131 homologs. The cytoplasmic TRAPID domain of human TMEM131 binds to TRAPPC8, which promotes collagen secretion as a key component of TRAPP III during the ER-to-Golgi transport of COPII vesicles (Fig. 7). Although we did not recognize any apparent homologous C-propeptide protein sequence in *C. elegans* COL-19 or COL-101, their normal secretion still requires TMEM-131 as *Drosophila* and human collagen secretions require TMEM-131 homologs. As COL-19::GFP secretion also requires *C. elegans* TRAPPC8 homolog, we propose that the interaction between TMEM131 and TRAPPC8 family proteins and their essential roles in collagen secretion are evolutionarily conserved, while human TMEM-131 evolved mechanisms to promote assembly of procollagens in the ER via direct binding of its PapD-L to procollagen. Together, these results have defined previously unknown physiological functions of conserved TMEM131 family proteins in collagen production and elucidated the underlying mechanism via TMEM131 interaction with PapD-L and TRAPPC8.

HSP47 is a collagen-specific chaperone that recognizes collagen trimers in the ER and prevents their premature aggregation during secretion (*48*). Collagen secretion also requires TANGO1, a protein that facilitates the assembly of a collagen export machine in the ER (*19, 20*). However, no apparent HSP47 and TANGO1 orthologs can be found in *C. elegans* or *Drosophila* (*21–23*). Compared with HSP47 or TANGO1, TMEM131 family proteins are evolutionarily more ancient, consistent with their broad requirements for collagen production in animals. In mammals, COL1A1/2 trimerization initiates from the C-propeptide domain, which can be replaced with a transmembrane domain without affecting trimer formation (*10*). This indicates that C-propeptide domains act by bringing procollagens close to ER membranes to facilitate procollagen assembly and secretion. Supporting this notion, we found direct COL1A2 binding to the ER-luminal TMEM131 PapD-L domains and striking collagen-production defects of TMEM131 deficient cells. Additional PapD-L interactors identified from Y2H screens suggest broader roles of TMEM131 beyond COL1A2 binding, although TMEM131 does appear to exhibit client-protein specificity as many other secreted proteins were not affected by LOF of *tmem-131* in *C. elegans* (Table S1, S2).

By yeast-two-hybrid assays, we found that TMEM131 PapD-L domains can interact with C-terminal pro-peptide domains of additional members of human collagen protein families (Fig. S7). With a similar intriguing function in secretion, bacterial PapD acts as a chaperone that recruits and assembles pilus components for cellular export through well-defined “donor-strand exchange” (DSE) mechanisms (*41, 49, 50*). DSE proceeds through a concerted beta strand displacement to orderly assemble pilus subunits before export to the bacterial periplasm. Whether PapD-L in TMEM131 acts by similar mechanisms for procollagens await further studies. Given our findings, we propose that TMEM131 family proteins play critical roles in recruiting procollagens on ER and/or secretory vesicular membranes to promote collagen assembly en route to secretion. As multicellular organisms evolved collagen-rich extracellular matrix (ECM), which is lacking in bacteria but analogous to bacterial pilus components, it is tempting to speculate that the role of PapD-L domains in collagen secretion originated from the role of PapD in secretion of bacterial pilus components.

Our study also raises the possibility of TMEM131 being a new therapeutic target for alleviating tissue fibrosis in human disorders, treatment of which remains a major unmet medical need (*9, 51*). Current anti-fibrotic clinical efforts mostly leverage our knowledge on the well-characterized pro-fibrotic mediators, including TGFß, which stimulates collagen gene expression and protein biosynthesis. As TGFß also controls gene expression involved in other biological processes than fibrosis, its inhibition can bring many side effects, including epithelial hyperplasia, abnormal immune and wound healing responses (*52*). Given the newly discovered role of TMEM131 in collagen secretion, inhibition of TMEM131 or its interfaces with collagen C-propeptide and TRAPPC8 may be therapeutically useful for treating pathological tissue fibrosis, including conditions in aging, systemic sclerosis, chronic inflammatory diseases, end-stage organ dysfunction and heart failure (*6, 9, 53, 54*).

## Acknowledgements

We thank the *Caenorhabditis* Genetics Center and National BioResource Project in Japan for *C. elegans* strains. The work was supported by NIH grants R01GM117461, R00HL116654, ADA grant 1-16-IBS-197, Hillblom foundation start-up grant, Pew Scholar Award, Klingenstein-Simons Fellowships in the Neurosciences, Alfred P. Sloan Foundation Fellowship, and Packard Fellowship in Science and Engineering (D.K.M). We thank Drs. Hong Zhang, Kamran Atabai, Akiko Hata and Matthew Shoulders for discussion and Dr. Jose Pastor-Pareja for the transgenic *Drosophila* line: w*; UAS-Cg25C.RFP.2.1 and Dr. Bruno Canque for TMEM131L plasmid reagents.

## Supplementary Materials

**Fig. S1.**
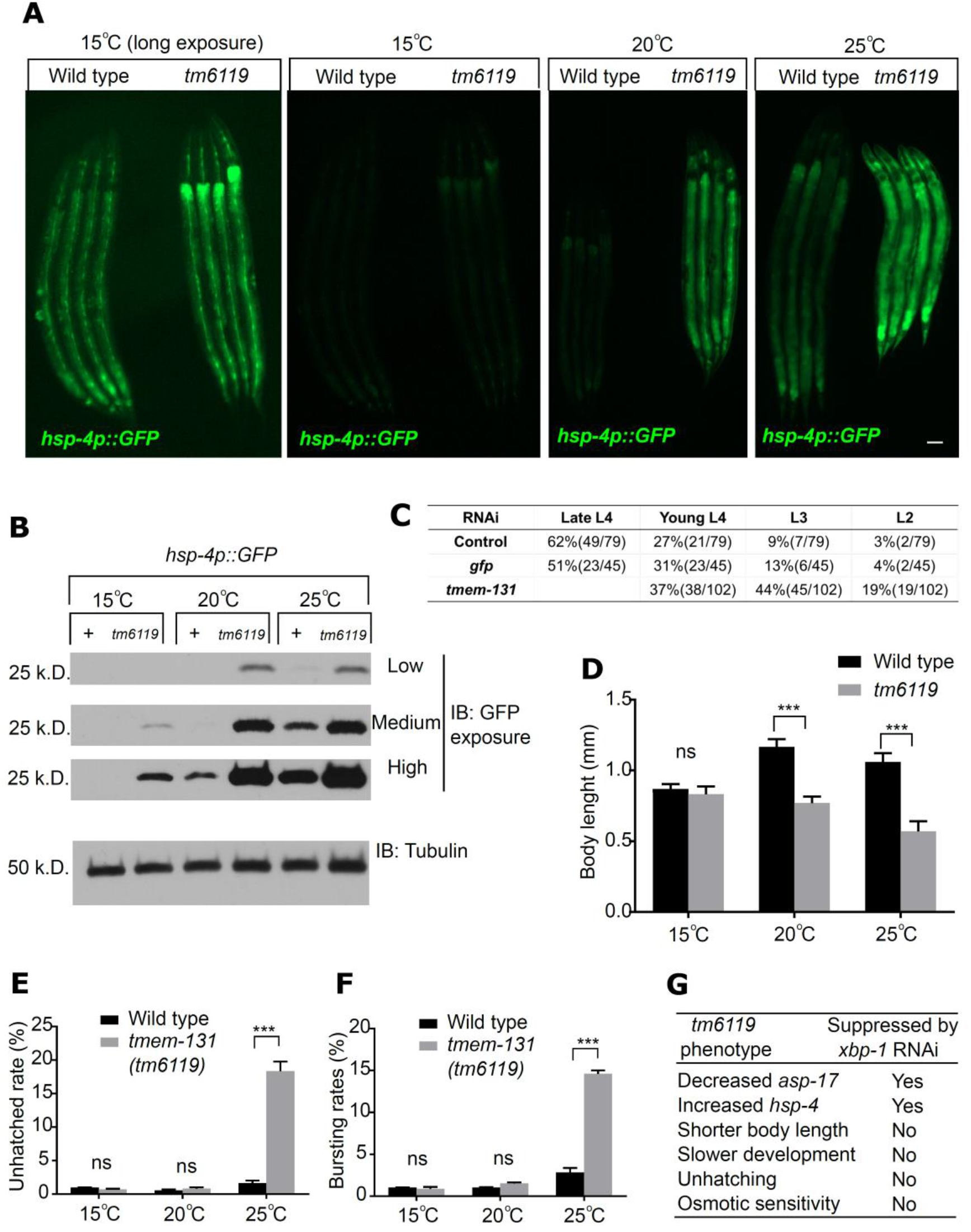
Additional phenotypic characterization of *tmem-131(tm6119)* mutants, Related to Fig. 1. **(A)**, Exemplar fluorescence images and (**B)**, Western blot analysis of *hsp-4p*::GFP protein abundance in wild type and *tmem-131(tm6119)* mutants under 15, 20 and 25 degrees. (**C)**, Quantification of developmental progression in animals with control, *gfp* and *tmem-131* RNAi in the presence of 1µg/mL ER stressor Tunicamycin. (**D-F)**, Quantification of animal body lengths (**D**), percentages of unhatched eggs (**E**) and osmotic stress-induced cuticle-bursting rates (**F**) in wild-type and *tmem-131(tm6119)* mutants under 15, 20 and 25 degrees. *** indicates P < 0.001 (n ≥ 3 biological replicates). n.s., no significant differences. Scale bars: 20 µm. **G**, Table summarizing the phenotype of *tmem-131(tm6119)* mutants and its dependency on XBP-1.

**Fig. S2.**
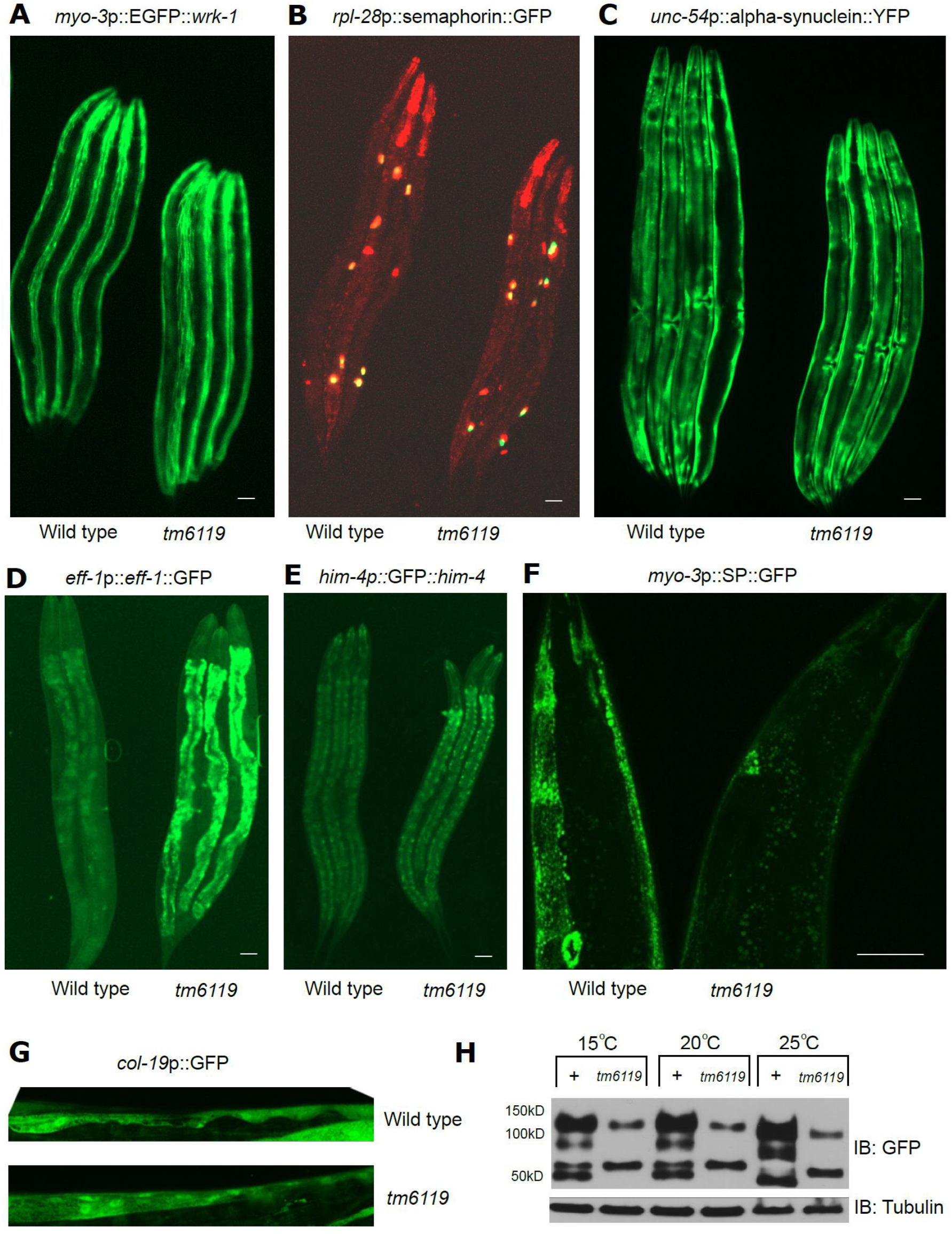
Fluorescent reporter screens and mutant verification for *tmem-131* phenotype, related to Fig. 2. **(A-E)**, Exemplar fluorescence images of translational reporters for (**A**) *wrk-1*, (**B**) semaphorin, (**C**) alpha-synuclein, (**D**) *eff-1,* (**E**) *him-4* and (**F**) *myo-2*p::ss (secretion peptide sequence)::GFP in wild-type and *tmem-131(tm6119)* mutants under 20 °C. (**G)**, Exemplar fluorescence images of *col-19* transcriptional reporter in wild type and *tmem-131*(*tm6119*) mutants under 20°C. (**H)**, Exemplar SDS-PAGE and Western blot analysis of COL-19::GFP in wild type and *tmem-131(tm6119)* mutants under 15, 20 and 25 degrees. Scale bars: 20 µm.

**Fig. S3.**
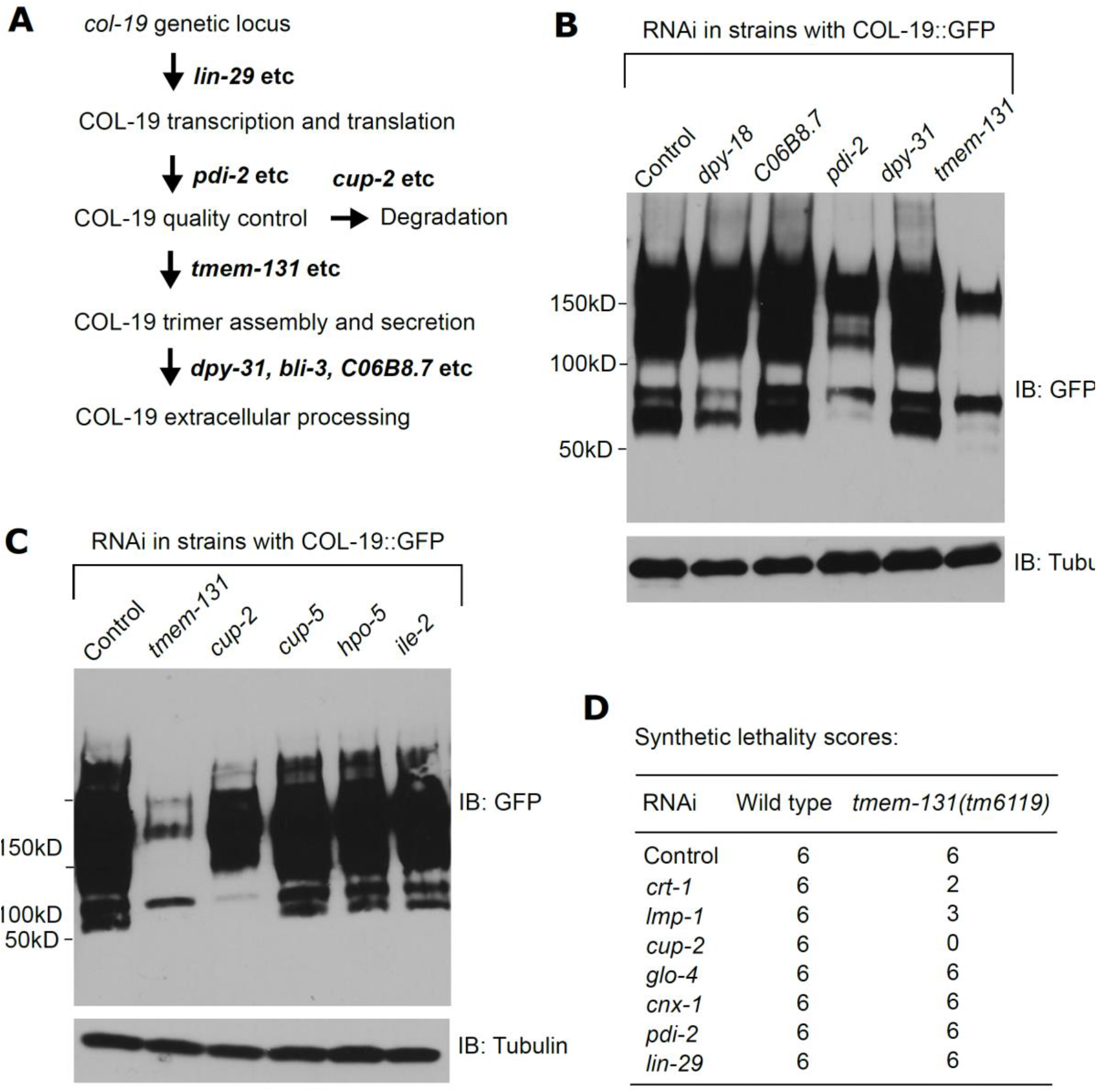
Genetic interaction of *tmem-131* with other genes involved in collagen production and degradation, Related to Fig. 3. **(A)**, Schematic diagram of the pathway and key gene regulators of COL-19 production and degradation in *C. elegans.* Place of *tmem-131* in the pathway is supported by evidence that *pdi-2* RNAi caused similar but slightly less severe phenotype as *tmem-131* and that *cup-2* RNAi caused strong and fully penetrant synthetic lethality with *tm6119* (B-D). (B), Exemplar SDS-PAGE and Western blot analysis of COL-19::GFP with RNAi knock-down of genes, including *pdi-2*, involved in collagen modification, folding or quality control pathways. (**C)**, Exemplar SDS-PAGE and Western blot analysis of COL-19::GFP with RNAi knock-down of genes, including *cup-2*, involved in collagen degradation-related lysosomal or ERAD pathways. (**D)**, Synthetic lethality test for *tmem-131* and genes involved in collagen degradation or processing. Growth scores from 0–6 were noted (0, 2 parental worms; 1, 1–10 progeny; 2, 11–50 progeny; 3, 51–100 progeny; 4, 101–150 progeny; 5, 151-200 progeny; and 6, 200+ progeny).

**Fig. S4.**
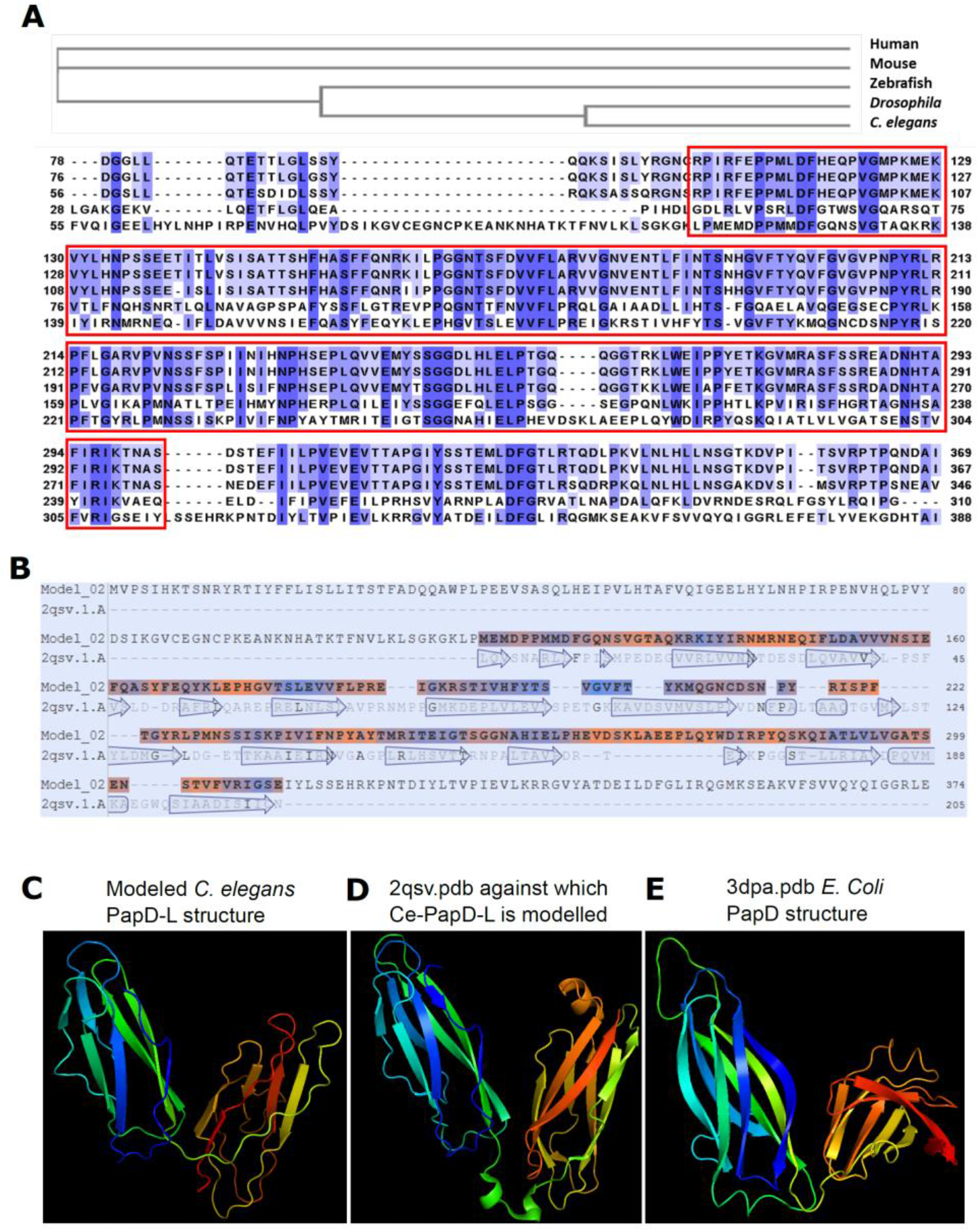
Sequence and structural similarities of Ce-PapD-L and other known PapD domain-containing proteins, Related Fig. 4. **(A)**, Multiple sequence alignment of conserved PapD-L domains from major metazoan species. (**B**), SWISS-MODEL comparison of PapD-L with bacterial PapD-L chaperone. (**C-E)**, Structures of *C. elegans* PapD-L (C, modelled against 2qsv.pdb, one of the most closely related structural homologs by SWISS-MODEL), 2qsv.pdb (**D**) and 3dpa.pdb *Ecoli*. PapD (**E**).

**Fig. S5.**
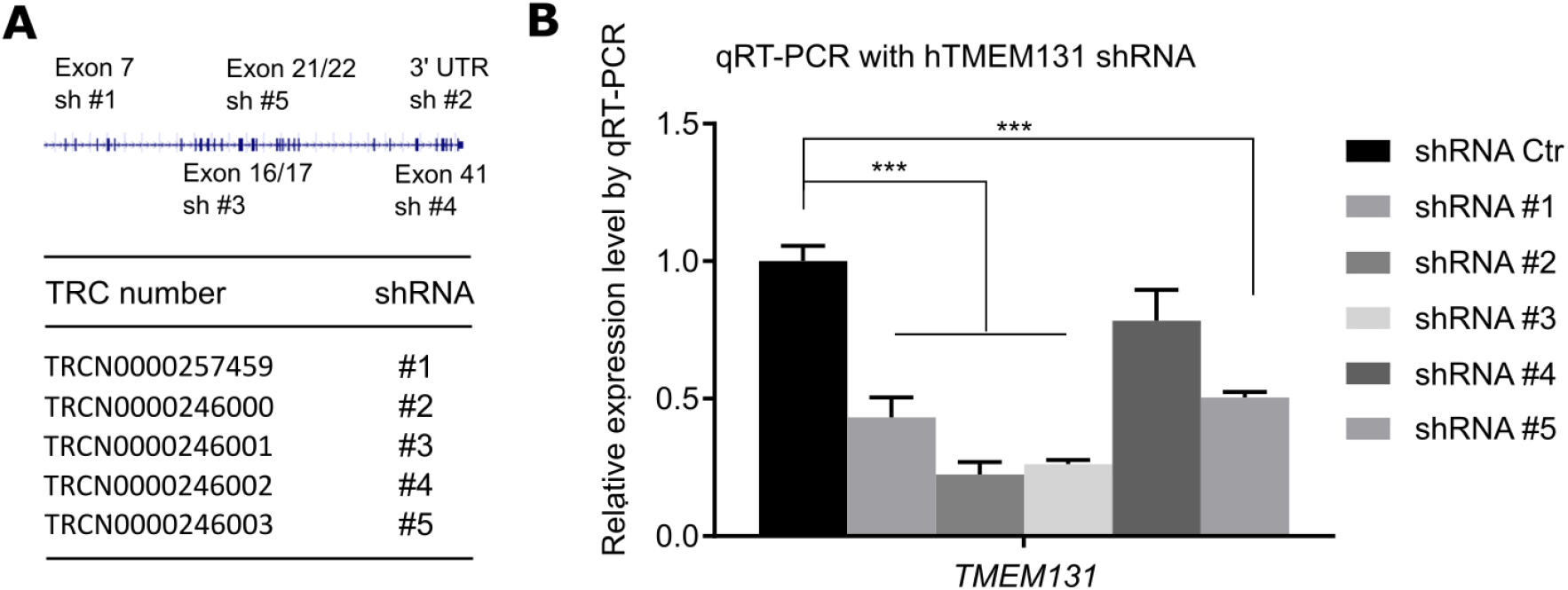
Verification of human *TMEM131* shRNA knockdown efficiency, Related to Fig. 4. **(A)**, Schematic illustrating 5 different shRNA sequences (from TRC consortium) targeting various coding sequences and 3’ UTR of human *TMEM131*. (**B)**, Quantitative RT-PCR measurements of human *TMEM131* expression levels in U2OS with lentiviral expression of indicated shRNAs.

**Fig. S6.**
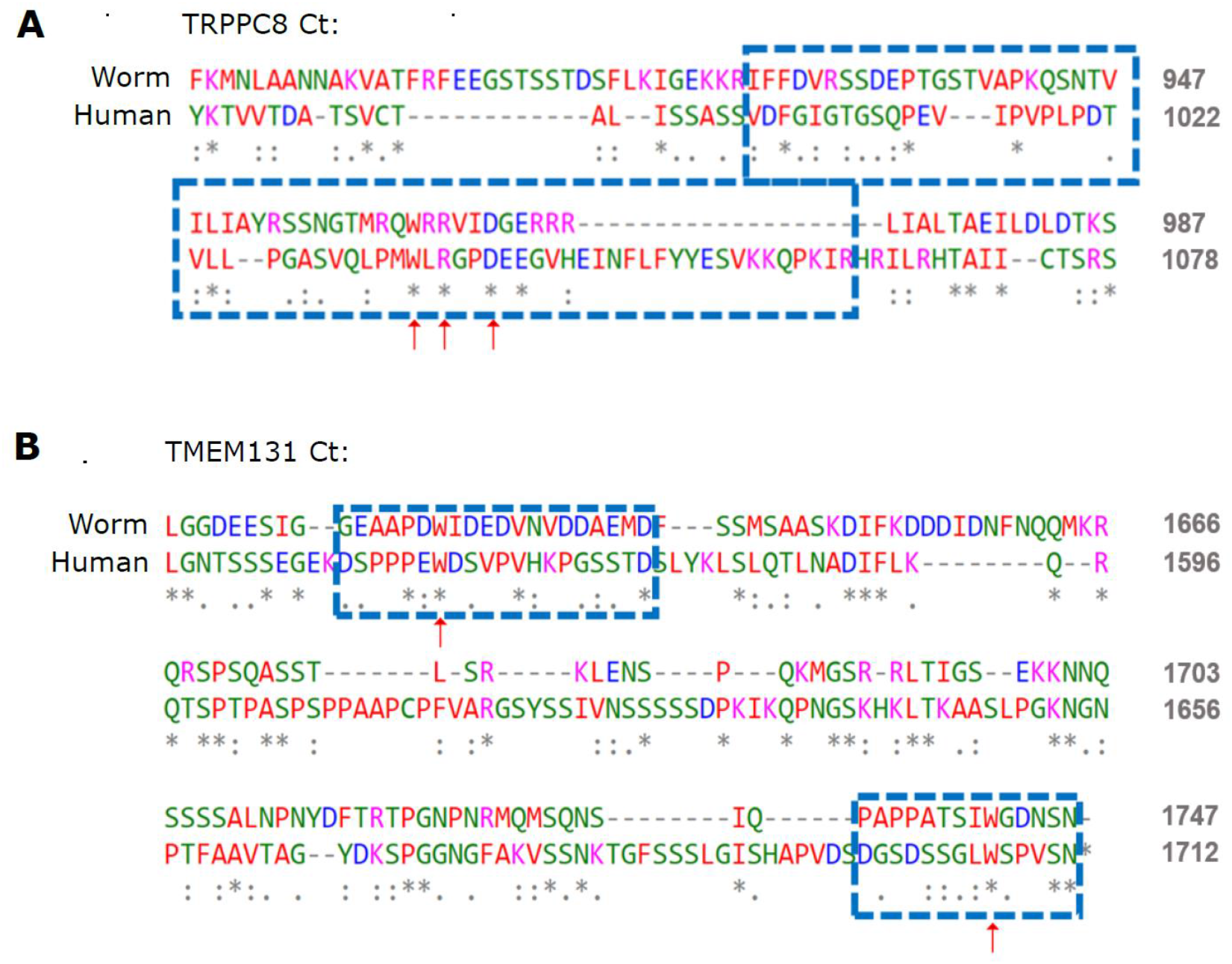
Conserved residues in TRAPPC8 and TMEM131 C termini, related to Fig. 6. (A), Sequence alignment of *C. elegans* and human TRAPPC8 C-termini indicating conserved WRD residues tested for importance in interaction with TMEM131. (B), Sequence alignment of *C. elegans* and human TMEM131 C-termini indicating conserved tryptophan residues tested for importance in interaction with TRAPPC8.

**Fig. S7.**
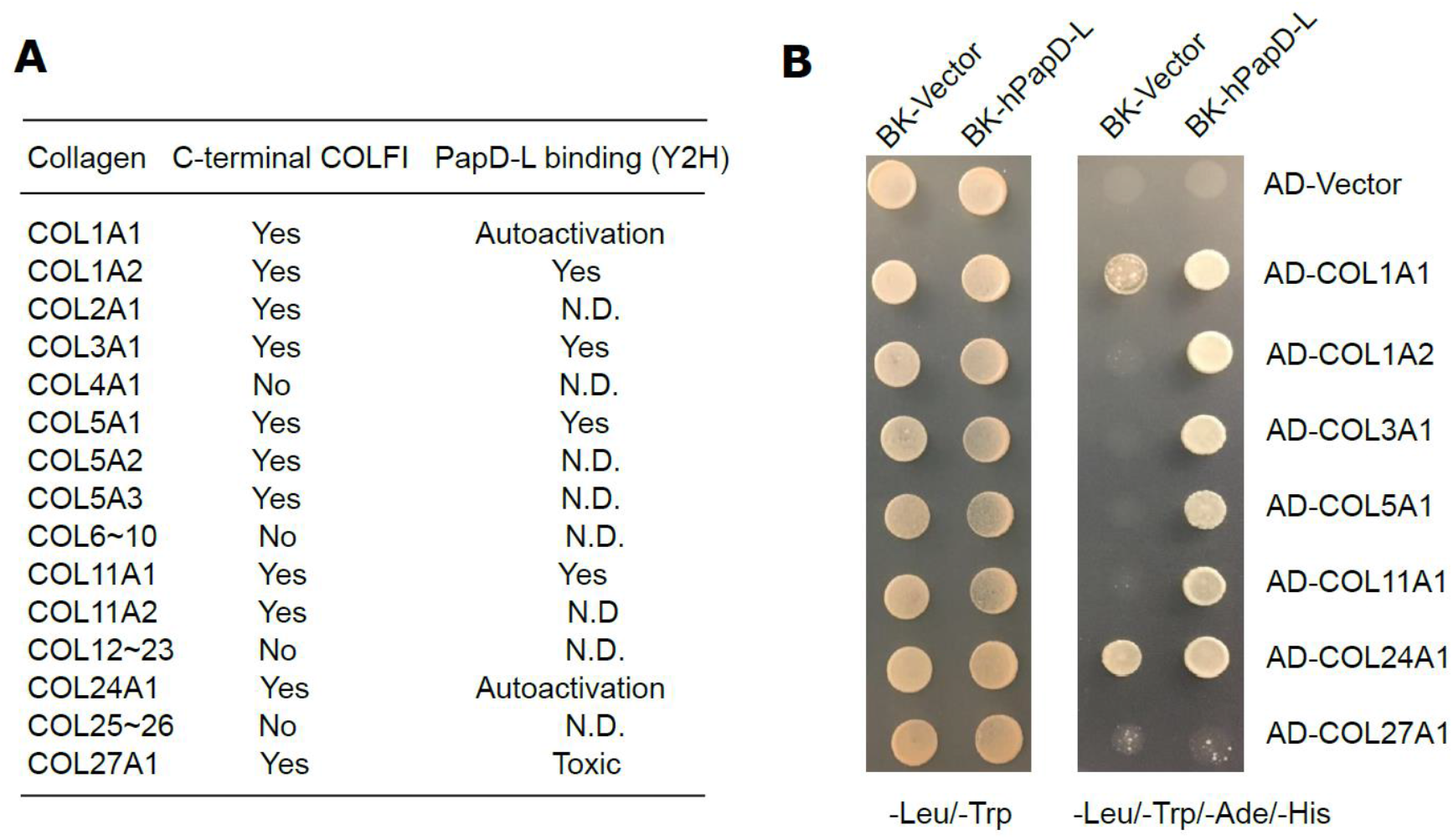
Y2H assay of interaction between PapD-L from TMEM131 and collagen family proteins with C-terminal COLFI domains, related to Fig. 7. **(A)**, Table summary of Y2H results showing interaction between PapD-L from human TMEM131 and specific collagen proteins. N.D., not determined owing to clone unavailability or little expression in yeasts. (**B)**, Exemplar yeast colony growth on indicated a.a. selection plates after transformation of plasmids encoding indicated bait and prey proteins for test of interaction in yeasts.

**Table S1. List of genes identified from genome-wide RNAi screen of *asp-17*p::GFP regulators.** (See Excel spreadsheet uploaded separately)

**Table S2.** Reporters examined in phenotypic screens for *tmem-131* RNAi.

| Genotype | Reporter | Control | *tmem-131* |
| --- | --- | --- | --- |
| *hsp-4p::GFP* | *hsp-4* transcriptional | * | **** |
| *hsp-16p::GFP* | *hsp-16* transcriptional | ** | ** |
| *col-19p::GFP* | *col-19* transcriptional | **** | **** |
| *col-19p::col-19::GFP* | *col-19* translational | **** | * |
| *col-101p::col-101::GFP* | *col-101* translational | **** | * |
| *cat-1p::cat-1::GFP* | *cat-1* C- terminal translational | ** | ** |
| *eff-1p::eff-1::GFP* | *eff-1* translational | * | *** |
| *emb-9p::emb-9::mCherry* | *emb-9* translational | *** | *** |
| *exl-1p::exl-1::GFP* | *exl-1* translational | *** | *** |
| *lrp-1p::lrp-1::GFP* | *lrp-1* translational | ** | ** |
| *fat-7p::fat-7::GFP* | *fat-7* translational | *** | *** |
| *gna-1p::GFP::gna-1* | *gna-1* translational | ** | ** |
| *him-4p::GFP::him-4* | *him-4* translational | ** | *** |
| *hmr-1p::hmr-1::GFP* | *hmr-1* translational | *** | * |
| *lam-1p::lam-1::dendra* | laminin-dendra translational | *** | *** |
| *lgg-1p::mCherry::GFP::lgg-1* | *lgg-1* translational | ** | ** |
| *lim-7p::ced-1::GFP* | *ced-1* translational | ** | ** |
| *mng::3xflag::cpg-1* | *cpg-1* translational | ** | ** |
| *myo-3p::EGFP::wrk-1* | *wrk-1* translational | *** | **** |
| *myo-3p::YFP::TRAM* | TRAM translational | ** | *** |
| *nhx-2p::cpl-1::YFP* | *cpl-1* translational | ** | *** |
| *nhx-2p::cpl-1(W32A Y35A)::YFP* | *cpl-1(W32A Y35A)* translational | *** | *** |
| *nhx-2p::ubiquitin-V::mCherry* | ubiquitin-V proteins | *** | *** |
| *rpl-28p::semaphorin::mCherry* | *F23H12.5* translational | **** | **** |
| *rpl-28p::manf-1::venus* | *manf-1* translational | **** | **** |
| *rpl-28p::T19D2.1::mCherry* | *T19D2.1* translational | **** | **** |
| *rpl-28p::Y73E7A.8::mCherry* | *Y73E7A.8* translational | **** | **** |
| *spon-1p::spon-1::vGFP* | *spon-1* translational | ** | ** |
| *unc-22p::egl-20::GFP* | *egl-20* translational | * | *** |
| *unc-54p::alpha-synuclein::YFP* | alpha-synuclein translational | **** | **** |
| *unc-54p::RFP::SP12* | ER membrane RFP marker | ** | ** |
| *unc-54p::mig-23::GFP* | *mig-23* translational | *** | *** |
| *vha-6p::mans::GFP* | intestinal GFP marker for the Golgi | *** | *** |
| *vit-2p::vit-2::GFP* | *vit-2* translational | * | *** |

\* indicates fluorescent reporter levels under control and *tmem-131* RNAi conditions;
qualitative changes were followed up by quantitative fluorescence and Western blot for verification.

**Table S3.** List of human cDNA clones identified by Y2H screens using Ce-PapD-L as a bait. Normalized human cDNA prey library were used for the screen. COL1A2 and WDR26 have been retested in Y2H assays for true interaction with Ce-PapD-L.

| Protein ID | Description |
| --- | --- |
| COL1A2 | Collagen alpha-2(I) chain |
| WDR26 | WD repeat domain 26 |
| FBLN5 | Fibulin 5 (secreted, extracellular matrix protein) |
| BICRAL | BRD4 interacting chromatin remodel complex associated protein |
| CGRRF1 | cell growth regulator with ring finger domain 1 |
| TGS1 | Trimethylguanosine synthase 1 |
| TKTL2 | Transketolase Like 2 |
| PON2 | Paraoxonase 2 |
| PCCB | Propionyl-CoA carboxylase enzyme |
| GPCPD1 | Glycerophosphocholine phosphodiesterase 1 |
| RNF10 | Rng finger protein 10 (ubiquitination) |
| MPP1 | Membrane palmitoylated protein 1 (guanylate kinase) |
| DYNC1I2 | Dynein cytoplasmic 1 intermediate chain 2 |
| SECISBP2 | SECIS binding protein 2 |
| HAX1 | Src family tyrosine kinases |
| ZNF655 | Misc_RNA |
| SNHG6 | Small nucleolar RNA host gene 6 |
| ZNF37BP | Zinc finger protein 37B, pseudogene |
| UBA5 | Ubiquitin like modifier activating enzyme 5 |
| PPIAP14 | Peptidylprolyl isomerase A pseudogene 14 |
| PNPT1 | Polyribonucleotide nucleotidyltransferase 1 |
| LPL | Lipoprotein lipase |
| GCA | Grancalcin |
| EMC4 | ER membrane protein complex subunit 4 |
| AMZ2P1 | Archaelysin family metallopeptidase 2 pseudogene 1 |
| ZNF280B | Zinc finger protein 280B |
| ATXN1 | A DNA-binding protein |
| KIA | Kizuna centrosomal protein |
| CPXM2 | Carboxypeptidase X, M14 family member 2 |
| OTULIN | OTU deubiquitinase with linear linkage specificity |
| MAPK10 | Mitogen-activated protein kinase 10 |
| VPS36 | Vacuolar Protein Sorting 36 |
| YWHAB | Tyrosine 3/tryptophan 5-monooxygenase activation protein beta |
| RGS5 | Regulator of G protein signaling 5 |

**Table S4.** List of human cDNA clones identified by Y2H screens using TMEM131 Ct as a bait. Normalized Universal human cDNA prey library was used for the screen.

| Protein ID | Description |
| --- | --- |
| C12orf4 | Mast cell degranulation, autosomal recessive intellectual disability |
| TM2D3 | TM2 domain containing 3, beta-amyloid binding protein |
| ARCN1 | Archain 1, Delta-COPI |
|  | Trafficking protein particle complex 8, ER-to-Golgi |
| TRAPPC8 | trafficking/COPII. |
| PIWIL1 | piwi like RNA-mediated gene silencing 1 |
| COMMD1 | copper metabolism domain containing 1 |
| RNF2 | E3 ubiquitin-protein ligase RING2 |
| CNRIP1 | cannabinoid receptor interacting protein 1 |
| CRISP2 | cysteine rich secretory protein, binds cholesterol, fibrils formation |
| ARL2BP | ADP ribosylation factor like GTPase 2 binding protein |
| ACADM | acyl-CoA dehydrogenase medium chain |
| EIF1B | translation initiation factor 1B |
| CPNE3 | Calcium-dependent phospholipid-binding protein |
| STX8 | syntaxin 8 |
| TKTL2 | transketolase like 2 |
| GPM6B | glycoprotein M6B |
| GFM2 | G elongation factor mitochondrial 2 |
| C1QTNF3 | Collagenous Repeat-Containing Sequence 26 KDa Protein |

**Table S5.** Genes required for COL-19::GFP production in *C. elegans* based on RNAi.

| Gene name | Description |
| --- | --- |
| *tmem-131* | Transmembrane protein 131 |
| *pdi-2* | Disulfide isomerase beta |
| *cup-2* | An ortholog of human DERL1 (derlin 1) |
| *sar-1* | Small guanosine triphosphatases |
| *sec-23* | Component of the coat protein complex II (COPII) |
| *rab-1* | Ortholog of human RAB1A and RAB1B |
| *uso-1* | USO1 vesicle transport factor |
| *trpp-3* | Trafficking protein particle complex 3 |
| *trpp-6* | Trafficking protein particle complex 6 |
| *trpp-8* | Trafficking protein particle complex 8 |

**Supplementary Table 6.**
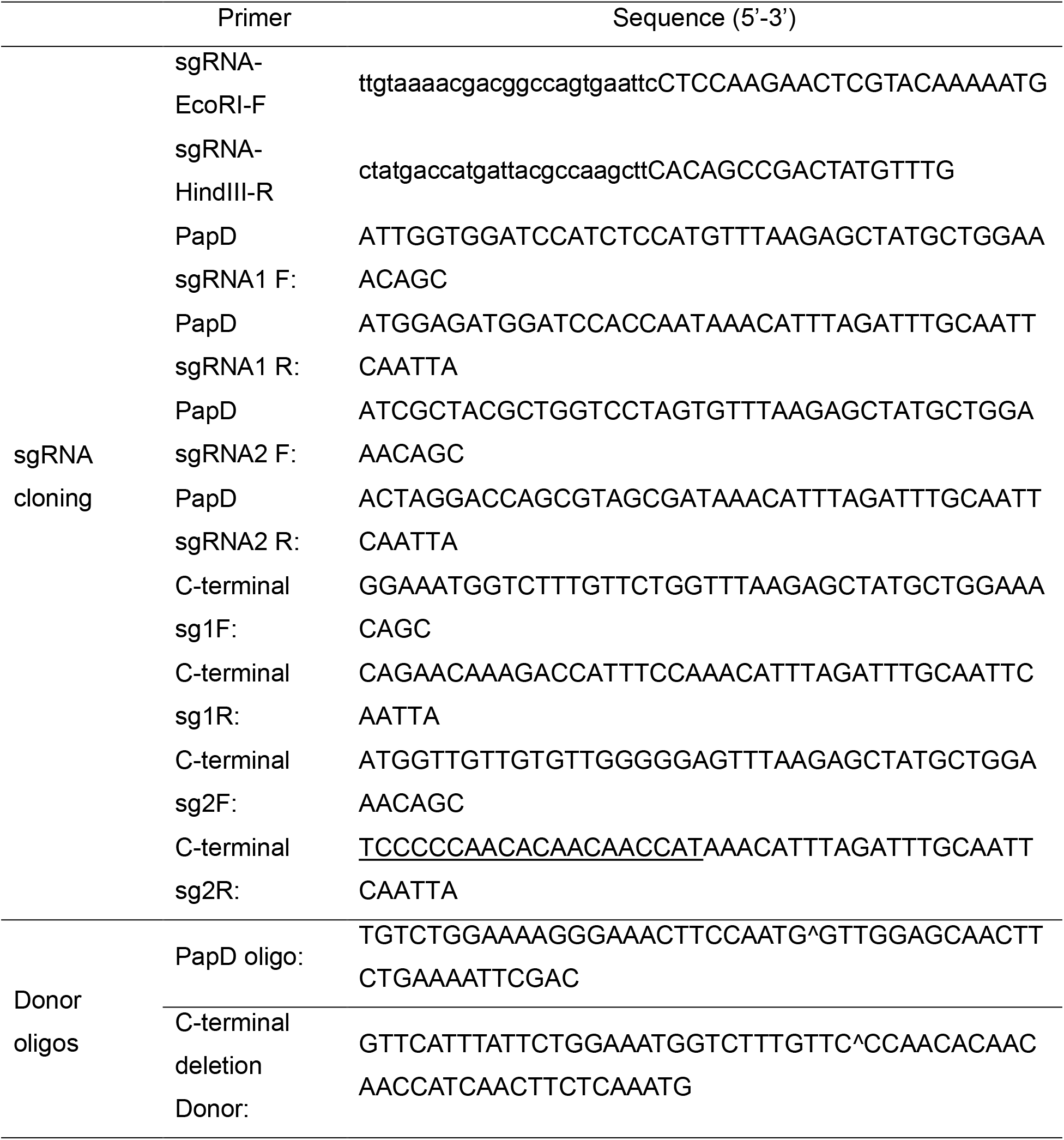
Primers and oligos used in genomic editing. Primers were used with Addgene plasmid #46169 as template. Primers with restriction sites were used for recombinant PCR with the respective PapD and C-terminal primers, which included a 20nt (bold and underlined) sgRNA target sequence from the *tmem-131* PapD and C-terminal region. The location of the deleted PapD and C-terminal motif is marked with “^”, generating a precise 1019 bp and 1730 bp deletion, respectively.

## Materials and Methods

### *C. elegans* culture and strains

*C. elegans* strains were grown on *E. coli* at 20°C using standard methods, unless otherwise specified (*55*). Synchronized worm populations were obtained by bleaching gravid adults. Feeding RNAi-mediated knock-down was performed as previously described (*56, 57*).

The N2 Bristol strain was used as the wild-type, and genotypes of strains used are as follows: *zcIs4* [*hsp-4*::GFP] V, *dmaIs8* [*hsp-16*p::GFP] IV, *rrf-3(pk1426)* II; *dmaIs10* [*asp-17*p::GFP, unc-54p::mCherry] X, *ire-1*(zc14) II; *zcIs4* V, *xbp-1*(*zc12*) III; *zcIs4* V, *kaIs12* [*col-19*::GFP], *dmaIs40* [*col-101*p::*col-101*::GFP; *unc-54*p::mCherry] and *tmem-131*(*tm6119*) III, which was further crossed with other reporters described above. Transgenic strains *dmaEx151* [*tmem-131*p::*tmem-131*::GFP; *myo-2*p::mCherry], *dmaEx146* [*tmem-131*p::GFP; *unc-54*p::mCherry], *dmaEx179* [*tmem-131*p::*tmem-131*::GFP; *myo-2*p::mCherry], *dmaEx152* [*rpl-28p::F23H12.5*::mCherry; *unc-122*p::GFP], *dmaEx153* [*rpl-28*p::*Y73E7A.8*::mCherry; *unc-122*p::GFP], *dmaEx169* [*rpl-28*p::*T19D2.1*::mCherry; *unc-122*p::GFP] were generated by germline transformation as described (*58*). The *tmem-131*(*dma301*) PapD-L deletion and *tmem-131(dma303)* C-terminal (1287–1808) deletion strains were generated by CRISPR/Cas9 methods to induce double-stranded breaks and subsequent homologous repair (Primer sequences are listed in Table S6). Other translational reporters used to identify a phenotype affected by RNAi against *tmem-131* and cross with *tmem-131*(*tm6119*) include *arIs37 [myo-3p::ssGFP + dpy-20(+)], bcIs39 [lim-7p::ced-1::GFP+lin-15(+)], rme-4(b1001); bIs1 [vit-2p::vit-2::GFP + rol-6(+)], caIs618 [eff-1p::eff-1::gfp], cpg-1(tn1728[mng::3xflag::cpg-1]) III, dmaEx115 [rpl-28p::manf-1::venus], dnSi4 [gna-1p::GFP + Cbr-unc-119(+)], juEx1111 [spon-1::vGFP], lrp-1(ku156)eqIs1 [lrp-1p::lrp-1::GFP]I; rrf-3(pk1426) II, muIs49 [egl-20::GFP+unc-22(+)], nIs590 [fat-7p::fat-7cod::GFP], nuIs26 [cat-1::GFP], pkIs2386 [unc-54p::alpha synuclein::YFP+ Cbr-unc-119(+)] IV;otIs181 [dat-1::mCherry + ttx-3::mCherry] III, otEx1184 [exl-1p::exl-1::GFP + rol-6(+)], osIs60 [unc-54p::MIG-23::GFP; unc-119(+)], osIs64 [myo-3p::YFP::TRAM;unc-119(+)], osIs66 [myo-3p:: EGFP::WRK-1], osIs77 [unc-54p::RFP::SP12; unc-119(+)], pwIs503 [vha-6p::mans::GFP+Cbr-unc-119(+)], qyIs44 [emb-9p::EMB-9::mCherry], qyIs108 [lam-1p::lam-1::dendra + unc-119(+)], rhIs23 III[GFP::him-4], sqIs11 [lgg-1p::mCherry::GFP::lgg-1 + rol-6(+)], veIs13 [col-19::GFP + rol-6(+)] V; let-7(mn112) unc-3(e151) X; mgEx725 [lin-4::let-7 + ttx-3::RFP], vkEx1243 [nhx-2p::ubiquitin-V::mCherry + myo-2p::GFP], vkEx1256 [nhx-2p::cpl-1::YFP + nhx-2p::dsRed::KDEL], vkEx1258 [nhx-2p::cpl-1(W32AY35A)::YFP + nhx-2p::DsRed::KDEL], vkEx1260 [nhx-2p::cpl-1::YFP + myo-2p::mCherry], vkEx1879 [nhx-2p::cpl-1(W32A Y35A)::YFP + myo-2p::mCherry], xnIs96 [hmr-1p::hmr-1::GFP]*. Extrachromosomal arrays were integrated using UV-irradiation and backcrossed for 3-6 times.

### Drosophila melanogaster experiments

Flies: UAS-Cg25C:RFP.2.1/CyO; Lsp2-Gal4/TM6B, and UAS-CG8370_dsRNA (Vienna Drosophila Resource Center ID# 42509/GD). Lsp2-Gal4 expresses specifically in the fat body. Flies expressing Collagen:RFP in fat body were crossed to either wild type or UAS-CG8370_dsRNA flies. Fat body was dissected from wandering stage third instar larvae and fixed in PFA 4%, stained with DAPI, and mounted for imaging by confocal microscopy.

### Quantitative RT-PCR

Total RNA was extracted following the instruction of Quick-RNA MiniPrep kit (Zymo Research, R1055), and reverse transcribed into cDNA (BioTools, B24408). Real-time PCR was performed by using SYBR Green Supermix (Thermo Fisher Scientific, FERK1081) on Roche LightCycler96 (Roche, 05815916001) system. Ct values of specific genes were normalized to measurements of *act-1* (*C. elegans*) and *RPL13* (human cell lines) levels. Results are presented as fold changes to respective references. Statistical significance was determined with t-test or two-way ANOVA, using GraphPad Prism 7. Primer sequences are listed in Supplementary Table 7.

### Imaging and fluorescence quantification

SPE confocal (Leica) and digital automated epifluorescence microscopes (EVOS, Life Technologies) were used to capture fluorescence images. Animals were randomly picked at the same stage and treated with 10 mM sodium azide and 1 mM levamisole in M9 solution (31742-250MG, Sigma-Aldrich), aligned on an 4% agar pad on a slide for imaging. Identical setting and conditions were used to compare experimental groups with control. For quantification of GFP fluorescence, animals were outlined and quantified by measuring gray values using the Image J software. The data were plotted and analyzed by using GraphPad Prism7.

### Western blot analysis of proteins

Stage-synchronized animals for control and experiment groups were picked (N > 40) and lysed directly into 20 µL Laemmli Sample Buffer for western blot analysis. Proteins were resolved by 15% SDS-PAGE (Bio-Rad, 4561084) and transferred to a nitrocellulose membrane (Bio-Rad, 1620167). Proteins of interest were detected using antibodies against GFP (A02020, Abbkine), Tubulin (Sigma, T5168) and H3 (Abcam, ab1791).

For subcellular fractionation, 50 ml adult-stage animal pellets were washed with M9 buffer for 3 times and resuspended in 500 μL RIPA lysis buffer (Amresco, N653) with 10 mM PMSF and protease inhibitor cocktail (BioTools, B14002). Then pellet samples are disrupted by TissueRuptor (Motor unit ‘8’ for 1 min) and incubated for 45 min in 4°C cold room. The lysate was centrifuged for 15 mins at 13,000 rpm for 15 mins, and the supernatant was collected as the soluble part, and the pellet was resuspended in 500 μL RIPA lysis buffer with 10 mM PMSF and protease inhibitor cocktail as the insoluble part. 20 µL samples added with equal volume of 2X Laemmli Sample Buffer were subject to Western blot analysis as described above.

### Yeast two hybrid assay

The cDNA coding sequences of the PapD-L domain of *C. elegans* TMEM-131 and the C-terminal TRAPID domain of human TMEM131 were cloned into the pGBKT7 vector and screened with a normalized universal human cDNA library (Clontech, 630481), following instructions in the Matchmaker® Gold Yeast Two-Hybrid System (Clontech, 630489). Verification of positive colonies was achieved by co-transforming PapD-L domains from different species (in pGBKT7 Vector) and genes of interest (in pGADT7 Vector) following the instruction of YeastMaker™ Yeast Transformation System 2 (Clontech, 630439) as well as plasmids from re-cloned cDNA.

### Genome-wide RNAi screen in *C. elegans*

Genome-wide RNAi screen in *C. elegans* was carried out by using the Ahringer library in a 96-well plate format modified from published protocol(*56, 57*). Briefly, dsRNA-expressing bacteria were replicated from Ahringer library to black-well, clear and flat bottom 96-well plate containing 100 µl of LB medium with 100 µg/ml Carbenicillin, and cultured overnight at 37 °C without shaking. 100 µl of LB containing 2-4 µg/mL of IPTG was added into each well to induce double-stranded RNA (dsRNA) expression for 2 hours. Bacteria containing dsRNA were then collected by centrifugation and re-suspended in 50 µl nematode growth (NG) medium containing 50 µg/ml Carbenicillin,1 µg/mL of IPTG and ∼10 synchronized L1 worms. The animals were cultured at 25 °C with shaking at the speed of 150 rpm for 3 days. Microscopic examination was then carried out looking for aberrant GFP-reporter expression and other phenotypes.

### Synthetic lethality analysis in *C. elegans*

Wild type and *tmem-131(tm6119)* mutants were grown on *E. coli* at 20 °C using standard methods for several (>2) generations. Two L3-L4 stages worms were picked into the RNAi plate for testing with the methods mentioned for Synthetic genetic-interaction (SGI) analysis, as described(*59*). *spg-7* (RNAi) was a positive control for RNAi efficiency. Over the course of 5 days, the number of progenies at adult stage were counted. The growth score was assigned from 0-6 (0, 2 parental worms; 1, 1-10 progeny; 2, 11-50 progeny; 3, 51-100 progeny; 4, 101-150 progeny; 5, 151-200 progeny; and 6, 200+ progeny), with at least three biological replicates.

### Lentiviral TMEM131 shRNA knockdown in cultured human HEK293 cells

Human TMEM131 knockdown in osteosarcoma U2OS cells was carried out with lentiviral shRNA (Sigma-Aldrich, SHCLNG-NM_015348). The TMEM131 shRNA targeting sequences are: 5’-TAGCAGTTTCTCACCTATAAT-3’(TRCN0000257459, CDS), 5’-ATTATGCGCCAAGATCTAATT-3’(TRCN0000246000, 3’UTR), 5’-TCCAATTGAGTTGGCTATAAA-3’(TRCN0000246001, CDS), 5’-CTCGGACCCTTGGTCTAATTC-3’(TRCN0000246002, CDS), 5’-CATAGATTGAGTGCTATATTT-3’ (TRCN0000246003, CDS). HEK293T was transfected by the pMD2.G, psPAX2 and shRNA plasmid, following the Lentivirus Production methods and manuals of TurboFect Transfection Reagent (Thermo Scientific, R0531). The lentivirus-based GFP-specific shRNA were used as negative controls (Addgene 31849). 48 hrs later, the TMEM131 shRNA lentivirus contain media was collected and filtrated by 0.45 µm syringe filter (Millipore EMD, SLHP033RS). The osteosarcoma U2OS cells were incubated with TMEM131 shRNA lentivirus medium for 24 hours in a humidified incubator at 37 °C with 5% CO2. Transduction efficacy was enhanced by adding Polybrene (Sigma-Aldrich TR-1003-G). Lentivirus-transduced cells were enriched by the medium with 1.5 μg/ml puromycin selection for 3 days. Ascorbates were exogenously supplemented to ensure proper collagen modification. The knockdown efficiency of TMEM131 shRNA was evaluated by qRT-PCR.

### Immuno-fluorescent staining of type I Procollagen in human U2OS cells

The human TMEM131 knockdown U2OS stable cells were seeded in 24-well plates with cover glass for 2 days, each with three replicates (Fisher Scientific, 22293232). After 1xPBS washing for once, cells were treated by 4% formaldehyde solution for 10 mins. With 1xPBS washing for three times, cells were treated with 0.2% Triton X-100 in 1xPBS solution for 15 mins. Then cells were incubated in 5% BSA in 1xPBS solution for 1 hr at 4 °C after 1xPBS washing for three times. Monoclonal anti-human Procollagen type I C-Peptide (PIP) clone PC5-5 (Takara Bio USA, M011) incubated with the cell sample at 4 °C for 12 hrs, then incubated with goat anti-mouse IgG (H+L) Alexa Fluor 488 oligoclonal secondary antibody for 1 hr (Fisher Scientific, A-11001). Following 1xPBS washing for three times, the cover slide with cell samples was sealed on the microscope slide with Fluoroshield Mounting Medium with DAPI (Thermo Fisher Scientific, NC0200574). For quantification of GFP fluorescence, every cell in the captured images was outlined and quantified by measuring gray values of fluorescence intensity using the Image J software. The data were plotted and analyzed by using GraphPad Prism7.

### Statistical analysis

Data are presented as means ± S.D. with p values calculated by unpaired Student’s t-tests and one-way ANOVA. Data with non-normal distribution, including qRT-PCR and penetrance results, were assessed by nonparametric Mann-Whitney and Fisher’s exact tests, respectively.

